# *In vivo* multiplex imaging of dynamic neurochemical networks with designed far-red dopamine sensors

**DOI:** 10.1101/2024.12.22.629999

**Authors:** Yu Zheng, Ruyi Cai, Kui Wang, Junwei Zhang, Yizhou Zhuo, Hui Dong, Yuqi Zhang, Yifan Wang, Fei Deng, En Ji, Yiwen Cui, Shilin Fang, Xinxin Zhang, Kecheng Zhang, Jinxu Wang, Guochuan Li, Xiaolei Miao, Zhenghua Wang, Yuqing Yang, Shaochuang Li, Jonathan Grimm, Kai Johnsson, Eric Schreiter, Luke Lavis, Zhixing Chen, Yu Mu, Yulong Li

**Affiliations:** Peking-Tsinghua Center for Life Sciences, Academy for Advanced Interdisciplinary Studies, Beijing 100871, China; PKU-IDG/McGovern Institute for Brain Research, Beijing 100871, China; State Key Laboratory of Membrane Biology, Peking University School of Life Sciences, Beijing 100871, China; Institute of Neuroscience, State Key Laboratory of Neuroscience, Center for Excellence in Brain Science and Intelligence Technology, Chinese Academy of Sciences, Shanghai 200031, China; Institute of Molecular Medicine, Peking University College of Future Technology, Beijing 100871, China; Neuroscience Institute, New York University Langone Medical Center, New York 10016, USA; Department of Anesthesiology, Beijing Chaoyang Hospital, Capital Medical University, Beijing 100020, China; Janelia Research Campus, Howard Hughes Medical Institute, Ashburn, Virginia 20147, USA; Department of Chemical Biology, Max Planck Institute for Medical Research, Heidelberg 69120, Germany; University of Chinese Academy of Sciences, Beijing 100049, China; Institute of Molecular Physiology, Shenzhen Bay Laboratory, Shenzhen, Guangdong 518055, China; National Biomedical Imaging Center, Peking University, Beijing 100871, China

**Keywords:** neurochemical, dopamine, far-red, sensor, multiplex imaging

## Abstract

Neurochemical signals like dopamine (DA) play a crucial role in a variety of brain functions through intricate interactions with other neuromodulators and intracellular signaling pathways. However, studying these complex networks has been hindered by the challenge of detecting multiple neurochemicals *in vivo* simultaneously. To overcome this limitation, we developed a single-protein chemigenetic DA sensor, HaloDA1.0, which combines a cpHaloTag-chemical dye approach with the G protein-coupled receptor activation-based (GRAB) strategy, providing high sensitivity for DA, sub-second response kinetics, and an extensive spectral range from far-red to near-infrared. When used together with existing green and red fluorescent neuromodulator sensors, Ca^2+^ indicators, cAMP sensors, and optogenetic tools, HaloDA1.0 provides high versatility for multiplex imaging in cultured neurons, brain slices, and behaving animals, facilitating in-depth studies of dynamic neurochemical networks.

## INTRODUCTION

Neuromodulators play an essential role in shaping behavior, in which specific neurons integrate a variety of neuromodulatory inputs to finely tune neural circuits via intracellular signaling mechanisms and pathways(*1*, *2*). The monoamine dopamine (DA) plays significant roles in reward, learning, and movement(*3–5*); moreover, the multifaceted role of DA in physiology is intricately linked with its interactions with other neuromodulators, including acetylcholine (ACh), endocannabinoids (eCBs), and serotonin (5-HT)(*6*, *7*). For example, emerging evidence suggests that under specific conditions ACh modulates the axonal release of DA in the striatum(*8–10*). Furthermore, DA’s downstream actions require its interaction with DA receptors and subsequent signal transduction via cytosolic second messengers such as cAMP and Ca^2+^(*11*, *12*). Consequently, obtaining a comprehensive view of DA’s functions requires precise examination of its intricate interactions within neurochemical networks, including its complex relationship with the other neuromodulators, and intracellular signaling molecules with high spatial and temporal resolution.

Achieving this goal requires tools that can be used to simultaneously monitor various neurochemical signals, including multiple neuromodulators and/or a combination of neuromodulators and cytosolic signaling molecules *in vivo*. Previously, our group and others developed a series of genetically encoded DA sensors based on the G protein-coupled receptor (GPCR) activation-based (GRAB) strategy, which can be used to visualize DA dynamics *in vivo* with exceptionally high spatiotemporal resolution(*13–17*). However, despite their advantages, these fluorescent sensors are limited to the green and red spectrum(*18*), restricting their use to dual-color imaging and limiting our ability to simultaneously track a large number of neurochemical signals. This has led to the urgent need to extend the spectral range of neuromodulator sensors, particularly to include far-red and near-infrared (NIR) wavelengths (> 650 nm). However, engineering genetically encoded far-red/NIR sensors is challenging due to the relatively low brightness of existing far-red/NIR fluorescent proteins and the difficulty associated with obtaining suitable circularly permutated far-red/NIR fluorescent proteins(*19*, *20*).

Combining the dye-capture protein HaloTag(*21*) with rhodamine derivatives offers a promising alternative approach, providing a broad spectral range, high brightness, and high photostability(*22*). Similar to GFP—for which the chromophore is context-sensitive— rhodamine derivatives also reside in an equilibrium between the closed, non-fluorescent lactone (L) form and the open, fluorescent zwitterionic (Z) form, and this equilibrium is affected by the surrounding environment(*23*, *24*). Although this chemigenetic strategy has been used successfully to develop far-red/NIR Ca^2+^ and voltage sensors(*25*, *26*), these sensors’ performance *in vivo* has not been studied.

Here, we combined our GRAB strategy with chemigenetics in order to develop a far-red DA sensor called GRAB_HaloDA1.0_ (hereafter referred to as HaloDA1.0). We then used this new sensor to perform three-color imaging with high spatiotemporal precision in a variety of *in vitro* and *in vivo* applications, including cultured neurons, acute brain slices, and behaving animal models.

## RESULTS

### Development and *in vitro* characterization of a far-red DA sensor

We used the human D1 receptor (D1R) as the DA-sensing module due to its superior membrane trafficking properties compared to other DA receptors(*13*). We started by replacing the third intracellular loop (ICL3) in D1R with an optimized circularly permutated HaloTag protein (cpHaloTag) originally derived from the Ca^2+^ sensor HaloCaMP(*25*). To optimize this new DA sensor, the chimera variants were labeled with far-red dyes conjugated to a HaloTag ligand (HTL), which form a covalent bond with the cpHaloTag(*27*). We generated the DA sensor based on the hypothesis that upon binding its ligand, the receptor undergoes a conformational change that in turn drives a conformational change in cpHaloTag, thereby shifting the equilibrium of the conjugated dye from the non-fluorescent (L) state to the fluorescent (Z) state, resulting in an increase in fluorescence (Fig. 1A). We then systematically optimized the cpHaloTag insertion sites, linker sequences, and critical residues in both cpHaloTag and D1R (Fig. S1), primarily using Janelia Fluor 646 (JF646) as the far-red dye(*27*). In total, we screened more than 2000 variants, resulting in the variant with the highest response, which we call HaloDA1.0 (Fig. 1B). We also generated a DA-insensitive sensor (called HaloDAmut) to use as a negative control by mutating sites in the receptor’s ligand-binding pocket (Figs. 1B and S1A).

**Fig. 1.**
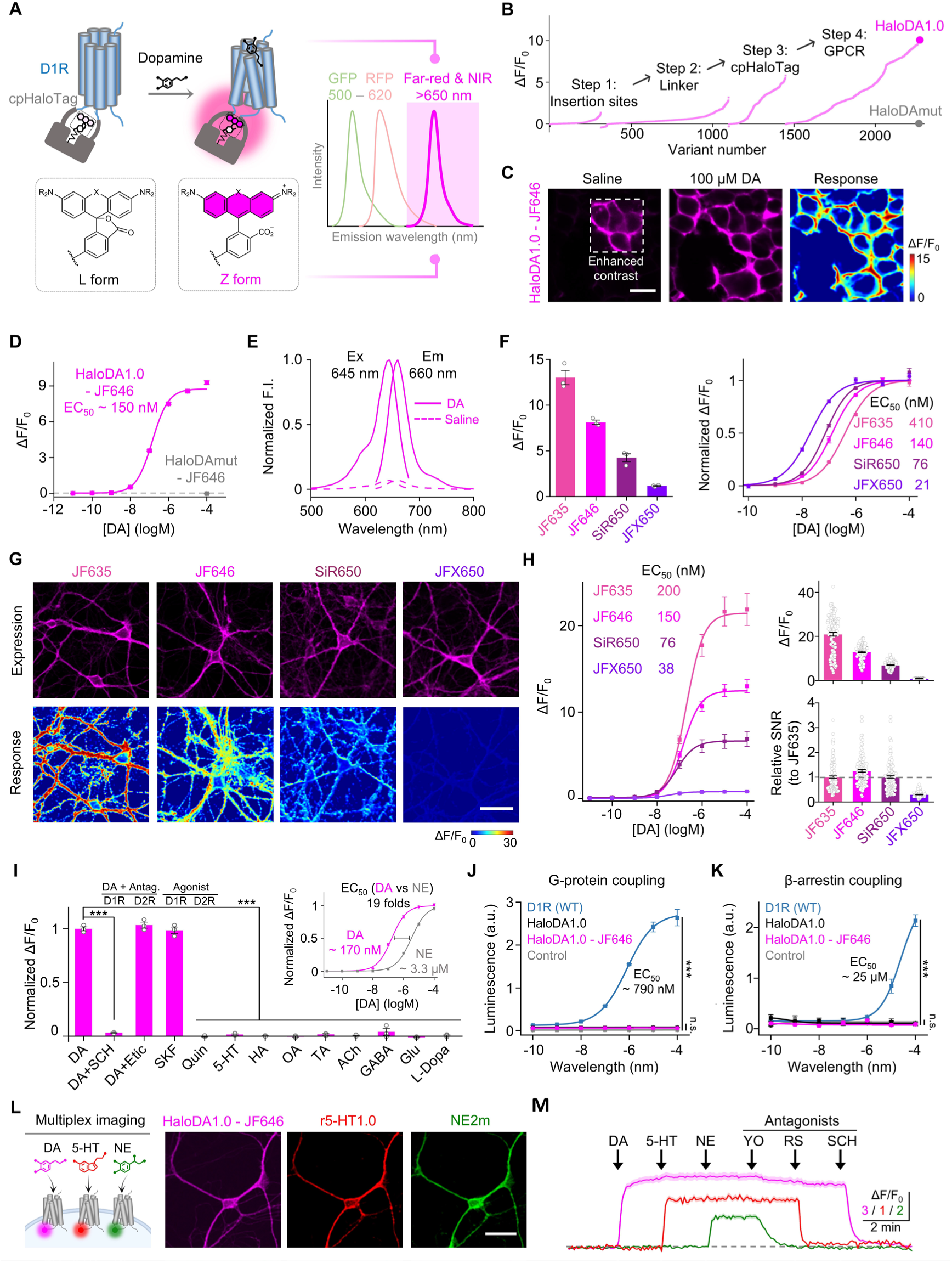
Development and characterization of a far-red dopamine sensor. (**A**) (Left) Schematic diagram illustrating the principle of the far-red dopamine (DA) sensor (top left) and L-Z equilibrium of rhodamine derivatives (bottom left). (Right) Idealized traces depicting the emission spectra of current GFP-and RFP-based sensors, alongside the new far-red and near-infrared (NIR) sensors. (**B**) Optimization of far-red DA sensor variants in response to 100 μM DA application, with stepwise changes in the insertion sites, linker, cpHaloTag and GPCR optimization. The variants in step 1 were screened using the dye JF635, while the variants in steps 2, 3, and 4 were screened using the dye JF646. (**C**) Representative images of HEK293T cells expressing HaloDA1.0 and labeled with JF646, before and after application of 100 μM DA. Scale bar, 20 μm. (**D**) Dose-response curves of HaloDA1.0 and HaloDAmut labeled with JF646 in HEK293T cells; n = 3 wells with 300–500 cells per well. (**E**) One-photon excitation (Ex) and emission (Em) spectra of HaloDA1.0 labeled with JF646 in the presence of 100 μM DA (solid lines) or saline (dashed lines). F.I., fluorescence intensity. (**F**) Maximum ΔF/F_0_ (left) and normalized dose-response curves (right) for HaloDA1.0 labeled with the indicated dyes in HEK293T cells; n = 3 wells with 300–500 cells per well for each dye. (**G**) Representative images of cultured rat cortical neurons expressing HaloDA1.0 and labeled with the indicated dyes (top row) and fluorescence response to 100 μM DA (bottom row). Scale bar, 50 μm. (**H**) Dose-response curves (left), maximum ΔF/F_0_ (top right), and signal-to-noise ratio (SNR) relative to JF635 (bottom right) for cultured rat cortical neurons expressing HaloDA1.0 and labeled with the indicated dyes; n = 120 regions of interest (ROIs) from 4 coverslips for each dye. (**I**) Normalized ΔF/F_0_ (relative to DA) for HaloDA1.0 expressed in cultured neurons and labeled with JF646. SCH, SCH-23390 (D1R antagonist); Etic, eticlopride (D2R antagonist); SKF, SKF-81297 (D1R agonist); Quin, quinpirole (D2R agonist); 5-HT, serotonin; HA, histamine; OA, octopamine; TA, tyramine; ACh, acetylcholine, GABA, γ-aminobutyric acid; Glu, glutamate; L-Dopa, levodopa. All chemicals were applied at 1 μM; n = 3 wells with an average of 50 neurons per well. The inset shows the dose-response curves for DA and norepinephrine (NE); n = 3–4 coverslips with 30 ROIs per coverslip. (**J**) Luciferase complementation assay to measure G protein coupling. Cells expressing miniGs-LgBit alone served as a negative control; n = 3 wells per group. WT, wild-type. (**K**) Tango assay to measure β-arrestin coupling. Non-transfected cells served as a negative control; n = 3 wells per group. (**L**) Schematic diagram depicting the strategy for multiplex imaging (left) and representative images (right) of cultured neurons co-expressing the far-red DA sensor (JF646-labeled HaloDA1.0), the red fluorescent 5-HT sensor (r5-HT1.0), and the green fluorescent NE sensor (NE2m). Scale bar, 50 μm. (**M**) Fluorescence responses of JF646-labeled HaloDA1.0 (magenta), r5-HT1.0 (red), and NE2m (green). Where indicated, DA (1 μM), 5-HT (1 μM), NE (1 μM), yohimbine (YO, 2 μM), RS23597-190 (20 μM), and SCH (10 μM) were applied; n = 40 ROIs from 3 coverslips.

We first confirmed that the JF646-conjugated HaloDA1.0 sensor (HaloDA1.0-JF646) traffics to the plasma membrane when expressed in HEK293T cells (Fig. 1C) and produces a strong, transient increase in fluorescence upon ligand application, with a half-maximal effective concentration (EC_50_) of 150 nM and a maximum ΔF/F_0_ of approximately 900% (Fig. 1D). Using one-photon excitation, we then confirmed that HaloDA1.0-JF646 is in the far-red range, with an excitation peak at 645 nm and an emission peak at 660 nm (Fig. 1E). Chemical dyes, which vary in their structure and properties, can affect the performance of HaloDA1.0; we therefore tested a wide range of rhodamine derivatives (*24*, *27–32*) in HaloDA1.0-expressing HEK293T cells, identifying several dyes that elicit a strong response in HaloDA1.0, with spectra spanning from green to NIR (Figs. 1F, S2, and S3). When labeled with distinct far-red dyes, HaloDA1.0 had peak ΔF/F_0_ responses ranging from 110% to 1300%, and EC_50_ values varying from 27 nM to 410 nM (Fig. 1F). Importantly, the DA-insensitive HaloDAmut sensor had no detectable fluorescence increase in response to DA application, regardless of the dye used (Figs. 1D and S2B). We also examined the performance of far-red dye–labeled HaloDA1.0 expressed in cultured neurons. Consistent with our results obtained with HEK293T cells, we observed a similar rank order for the four dyes tested in terms of the sensor’s peak response and DA affinity (Fig. 1G, H). Together, these results indicate that the properties of HaloDA1.0—including its spectrum, ligand response, and ligand affinity—can be fine-tuned by labeling with specific chemical dyes.

**Fig. 2.**
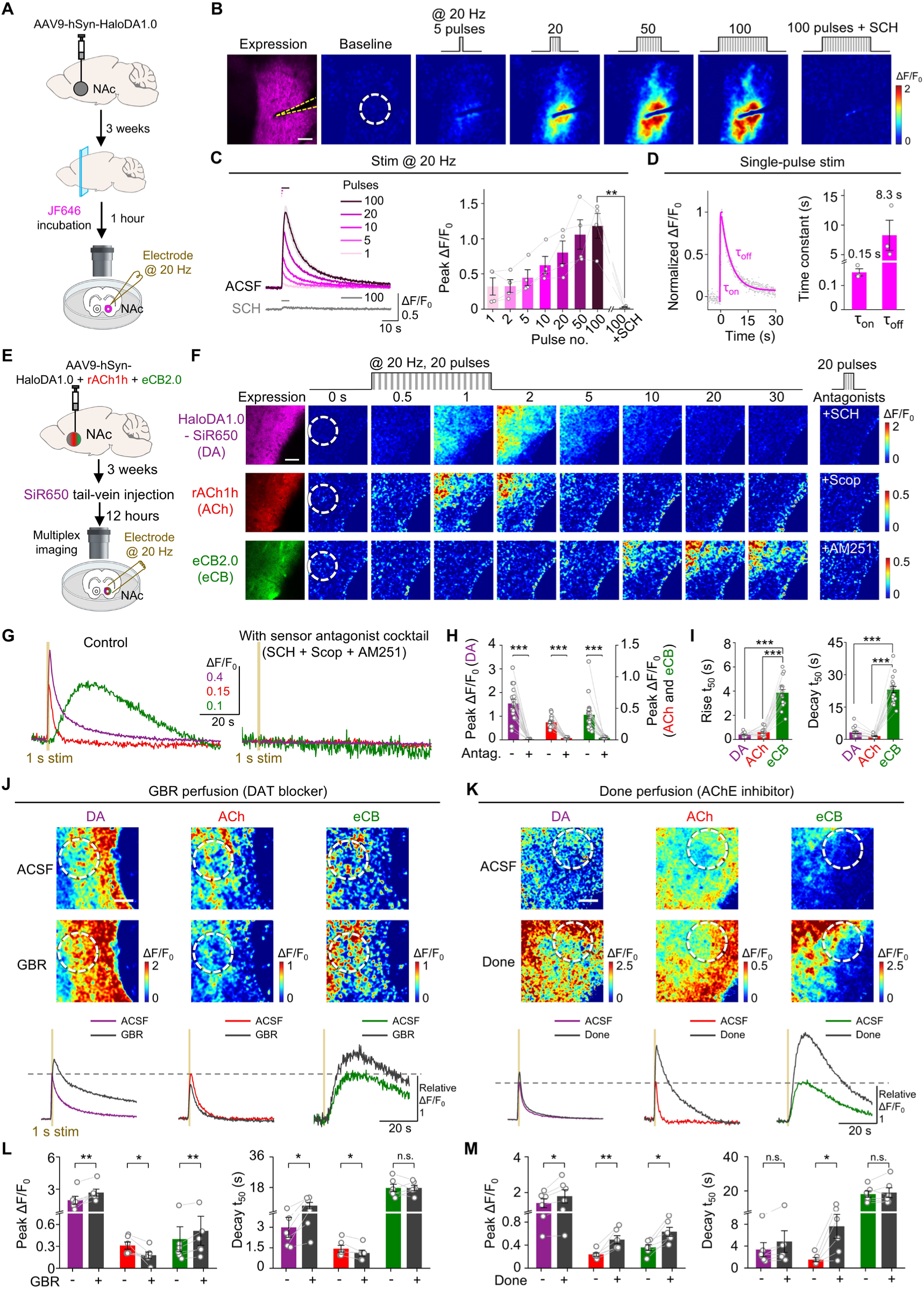
Multiplex imaging using HaloDA1.0 in acute brain slices. (**A**) Schematic illustration depicting the strategy for single-color imaging of mouse brain slices expressing HaloDA1.0. (**B**) Representative images showing the expression and fluorescence response of JF646- labeled HaloDA1.0 at baseline and in response to the indicated electrical stimuli. The white dashed circle (100 μm diameter) indicates the ROI used for further analysis, and the approximate location of the stimulating electrode is indicated with dashed yellow lines. SCH, 10 μM. Scale bar, 50 μm. (**C**) Representative traces (left) and group summary (right) of the changes in JF646-labeled HaloDA1.0 fluorescence in response to the indicated number of electrical stimuli; n = 4 slices from 3 mice. (**D**) Representative trace showing normalized ΔF/F_0_ (left) and group summary of τ_on_ and τ_off_ (right) measured in response to a single electrical stimulus. The trace was fitted with single-exponential functions to determine τ_on_ and τ_off_; n = 3 slices from 3 mice. (**E**) Schematic illustration depicting the strategy for multiplex imaging of mouse brain slices prepared 12 hours after injecting 100 nmol SiR650 into the mouse’s tail vein. (**F**) Representative images showing the expression and time-lapse fluorescence responses of SiR650-labeled HaloDA1.0, rACh1h, and eCB2.0 in response to the indicated electrical stimuli. The fluorescence response of each sensor measured in the presence of its corresponding antagonist (SCH, scopolamine, or AM251, applied at 10 μM) is shown on the far right. The white dashed circle (100 μm diameter) indicates the ROI used for further analysis. Scale bar, 50 μm. (**G** and **H**) Representative traces (**G**) and group summary (**H**) of the fluorescence change in SiR650-labeled HaloDA1.0 (magenta), rACh1h (red), and eCB2.0 (green) in response to electrical stimuli (20 Hz applied for 1 s) before and after application of the antagonist cocktail; n = 18 slices from 6 mice. (**I**) Group summary of the rise and decay kinetics (t_50_) of all three sensors in response to electrical stimuli. (**J** and **K**) Representative pseudocolor images (top) and traces of the fluorescence response (bottom, relative to ACSF) of the indicated sensors in response to electrical stimuli (20 Hz applied for 1 s) before and after the application of 2 μM GBR (**J**) or donepezil (**K**). Scale bar, 50 μm. (**L** and **M**) Group summary of peak ΔF/F_0_ (left) and decay t_50_ (right) for the indicated three sensors in response to electrical stimuli in the presence of GBR (**L**) or donepezil (**M**); n = 6 slices from 3 mice for each treatment.

Next, we characterized the sensor’s pharmacological properties, kinetics, and coupling to downstream pathways when expressed in HEK293T cells and cultured neurons and then labeled with either JF646 or SiR650(*28*) (Figs. 1I-K and S4). We found that HaloDA1.0 retains the pharmacological properties of the parent receptor, as it can be activated by the D1R agonist SKF-81297, but not the D2R-specific agonist quinpirole. In addition, the DA-induced increase in HaloDA1.0 fluorescence was blocked by co-application of the D1R-specific antagonist SCH-23390 (SCH), but was unaffected by the D2R-specific antagonist eticlopride (Figs. 1I and S4A). Moreover, HaloDA1.0 has 15-19-fold higher sensitivity to DA compared to the structurally similar neuromodulator norepinephrine (NE), and had only a minimal response to a wide range of other neurochemicals tested (Figs. 1I and S4A, B). Using line-scan confocal microscopy, we then locally applied DA followed by SCH in order to measure the sensor’s on-rate (τ_on_) and off-rate (τ_off_), respectively. We found that HaloDA1.0 has a sub-second on-rate (with τ_on_ values of 40 ms and 90 ms when labeled with JF646 and SiR650, respectively), and an off-rate similar to values reported for DA sensors (with τ_off_ values of 3.08 s and 2.96 s when labeled with JF646 and SiR650, respectively) (Fig. S4C, D). To examine whether HaloDA1.0 couples to downstream intracellular signaling pathways, we used the luciferase complementation assay and the Tango assay to measure Gs-and β-arrestin–mediated signaling, respectively. Importantly, we found that HaloDA1.0 induces only minimal activation of these two signaling pathways (Fig. 1J, K); as a positive control, we found that the wild-type D1R has robust dose-dependent coupling to both pathways (Fig. 1J, K). As an additional verification, we examined whether HaloDA1.0 undergoes β-arrestin–mediated internalization and/or desensitization when expressed in cultured neurons. We found that the DA-induced increase in HaloDA1.0 surface fluorescence was stable for at least 2 hours, indicating minimal internalization (Fig. S4E, F). Taken together, these results indicate that HaloDA1.0 has high sensitivity and specificity for DA, with rapid response kinetics, but without the complication of activating downstream signaling pathways.

The most significant advantage of our far-red HaloDA1.0 sensor is its potential for use in multiplex imaging when combined with green and/or red fluorescent sensors. As an initial proof of concept, we co-expressed the far-red HaloDA1.0-JF646 sensor, the red fluorescent 5-HT sensor r5-HT1.0(*33*), and the green fluorescent NE sensor NE2m(*34*) in cultured neurons and then performed three-color imaging using confocal microscopy. We found that all three sensors were expressed in the same neuron, and their respective fluorescence signals could be sequentially activated and blocked by application of their respective agonists and antagonists, allowing us to simultaneously monitor all three monoamine neuromodulators in real time, with minimal crosstalk (Fig. 1L, M).

### The HaloDA1.0 sensor is compatible for use in multiplex imaging in acute brain slices

To assess whether HaloDA1.0 can detect endogenous DA release, we injected an adeno-associated virus (AAV) expressing HaloDA1.0 into the nucleus accumbens (NAc) of mice. After three weeks (to allow for expression), we prepared acute brain slices and labeled the sensor by incubating the slices for 1 hour with JF646 (Fig. 2A). we found that applying local electrical stimuli at 20 Hz elicited a robust, transient increase in fluorescence, with the amplitude of the response correlated with the number of stimuli (Fig. 2B, C). The sensor’s specificity for endogenous DA release was confirmed by application of the D1R antagonist SCH, which completely blocked the fluorescence increase (Fig. 2B, C). Importantly, HaloDA1.0 is highly sensitive, as it was able to detect DA release induced by a single electrical pulse, with a mean rise time constant of 150 ms and a mean decay time constant of 8.3 s (Fig. 2D).

In addition to DA, a variety of other neuromodulators such as ACh and eCBs are also released in the NAc, and although their interaction with DA has physiological relevance, this crosstalk between neuromodulator systems remains poorly understood(*6*, *35*). This intricate network of neuromodulators in the NAc therefore provides an excellent model system for performing multiplex imaging in a physiological context. We injected a mixture of viruses expressing HaloDA1.0 (subsequently labeled with the far-red dye SiR650 by tail vein injection), rACh1h, and the green fluorescent eCB sensor eCB2.0(*36*) in the NAc, allowing us to monitor all three neuromodulators simultaneously using confocal microscopy in acute brain slices (Fig. 2E). We found that all three sensors had a robust increase in their respective fluorescence signals in response to field stimuli applied at 20 Hz; moreover, each sensor’s signal was blocked by application of its respective antagonist, confirming specificity (Fig. 2F-H). Interestingly, compared to the DA and ACh signals, the eCB signal had significantly slower rise and decay times, with an onset of eCB release occurring after the end of the stimulus (Fig. 2I). This difference in release kinetics between neuromodulators is presumably due to differences in their respective release mechanisms, as eCB must be synthesized before it can be released(*37*, *38*), while DA and ACh are directly released from preloaded vesicles upon stimulation(*39*, *40*).

Our successful use of three-color imaging to measure three distinct neuromodulators provides a good system in which to study their regulation. We therefore pre-treated brain slices with the selective DA transporter (DAT) blocker GBR192909 (GBR) and found that GBR both increased the peak response and slowed the decay kinetics of the stimulus-induced DA signal (Fig. 2J, L), consistent with reduced reuptake of DA into the presynaptic terminal. Moreover, GBR significantly reduced the peak ACh signal (Fig. 2J, L), consistent with the notion that DA inhibits ACh release by binding D2R on cholinergic interneurons(*41–43*); in addition, GBR significantly increased the peak eCB signal (Fig. 2J, L). Similarly, the acetylcholinesterase inhibitor donepezil increased the stimulus-induced ACh signal, and significantly—albeit modestly—increased the DA signal (Fig. 2K, M), consistent with ACh’s known mechanism of action via nicotinic ACh receptors at dopaminergic terminals(*8*, *44*, *45*). Interestingly, donepezil also increased the peak eCB signal (Fig. 2K, M), suggesting a previously unknown interaction between the ACh and eCB signaling pathways. Together, these results demonstrate that HaloDA1.0 is suitable for use in multiplex imaging and provide new insights into the crosstalk between three key neuromodulators.

### Multiplex imaging in zebrafish larvae

To examine whether HaloDA1.0 can be used to monitor DA dynamics *in vivo*, we transiently expressed the sensor in neurons in larval zebrafish, leveraging their genetic accessibility and optical transparency. We then labeled the sensor using three far-red dyes—JF635, JF646, and SiR650—and found that SiR650-labeled sensors had the strongest baseline fluorescence (Fig. S5A, C). Moreover, locally applying a puff of DA rapidly induced a robust, transient increase in fluorescence, with the largest response measured in JF646-labeled sensors (Fig. S5B, C). SiR650-and JF646-labeled sensors had a similar signal-to-noise ratio (SNR), significantly outperforming JF635-labeled sensors. As negative controls, we confirmed that a puff of phosphate-buffered saline (PBS) had no effect on SiR650-labeled HaloDA1.0, and DA had no effect on SiR650-labeled HaloDAmut.

We then performed three-color *in vivo* imaging in zebrafish larvae by transiently expressing HaloDA1.0 in a zebrafish line expressing the red fluorescent Ca^2+^ sensor jRGECO1a in neurons and the green fluorescent ATP sensor ATP1.0(*46*) in astrocytes (Fig. S6A); the HaloDA1.0 sensor was then labeled with SiR650. Upon application of a mild electrical body shock, we observed time-locked fluorescence increases for all three sensors in the hindbrain (Fig. S6B1, C1). The kinetics of the DA and ATP signals were similar, but both signals decayed more slowly than the neuronal Ca^2+^ signal (Fig. S6D). A correlation analysis confirmed the strong correlation between the DA and ATP signals, with a negligible time lag between these two signals (Fig. S6D). In addition, application of the GABA_A_ receptor antagonist pentylenetetrazole (PTZ) induced robust, synchronized DA and ATP signals that were in phase with the neuronal Ca^2+^ signal (Fig. S6B2, C2); by aligning the DA and ATP signals with the peak Ca^2+^ signal, we found a high correlation in peak amplitude between the DA and Ca^2+^ signals and between the ATP and Ca^2+^ signals (Fig. S6E). Interestingly, we found that the decay kinetics of the DA signals differed between signals induced by electrical shock and signals induced by PTZ application; in contrast, we found no difference in decay kinetics for the Ca^2+^ and ATP signals (Fig. S6F). Taken together, these data indicate that the HaloDA1.0 sensor can reliably detect DA release *in vivo* and is compatible for use in three-color imaging in the brain of zebrafish larvae.

### HaloDA1.0 can detect optogenetically evoked DA release in freely moving mice

Using a cpHaloTag-based sensor *in vivo* in mice requires delivery of the dye to the mouse’s brain, presenting a greater challenge compared to its use in zebrafish. Therefore, we systematically compared various far-red dyes *in vivo* in order to optimize the performance of HaloDA1.0. We virally expressed the optogenetic actuator ChR2 (Channelrhodopsin-2) in the ventral tegmental area (VTA), and we expressed HaloDA1.0 in the NAc (Figs. 3A and S7A), which receives dense dopaminergic projections from the VTA. We then injected various dyes into the tail vein (to label HaloDA1.0 in the NAc), and performed fiber photometry recordings 12 hours later. Optogenetic stimulation of the VTA resulted in a moderate increase in JF646-labeled HaloDA1.0 fluorescence, with no measurable change in JF635-labeled or JFX650-labeled HaloDA1.0 (Fig. 3A2-A4). In contrast—and consistent with our results obtained with zebrafish—SiR650-labeled HaloDA1.0 had a much higher response. As negative controls, no signal was detected in uninjected mice or in mice expressing SiR6560-labeled HaloDAmut (Fig. 3A2-A4). In addition, an intraperitoneal (i.p.) injection of the DAT blocker GBR produced a slow progressive increase in the basal fluorescence of SiR650-labeled HaloDA1.0 and increased both the magnitude and decay time of the light-activated responses (Fig. 3B). Moreover, the D1R antagonist SCH application abolished both the increase in basal fluorescence and the light-evoked responses. The optogenetically evoked signals were stable for two days but then decreased, presumably due to degradation of the sensor-dye complex, as the responses were restored by subsequent injections of dye (Fig. S8). These results indicate that expressing HaloDA1.0 and then labeling the sensor with SiR650 provides a sensitive and specific tool for monitoring the release of endogenous DA *in vivo*.

**Fig. 3.**
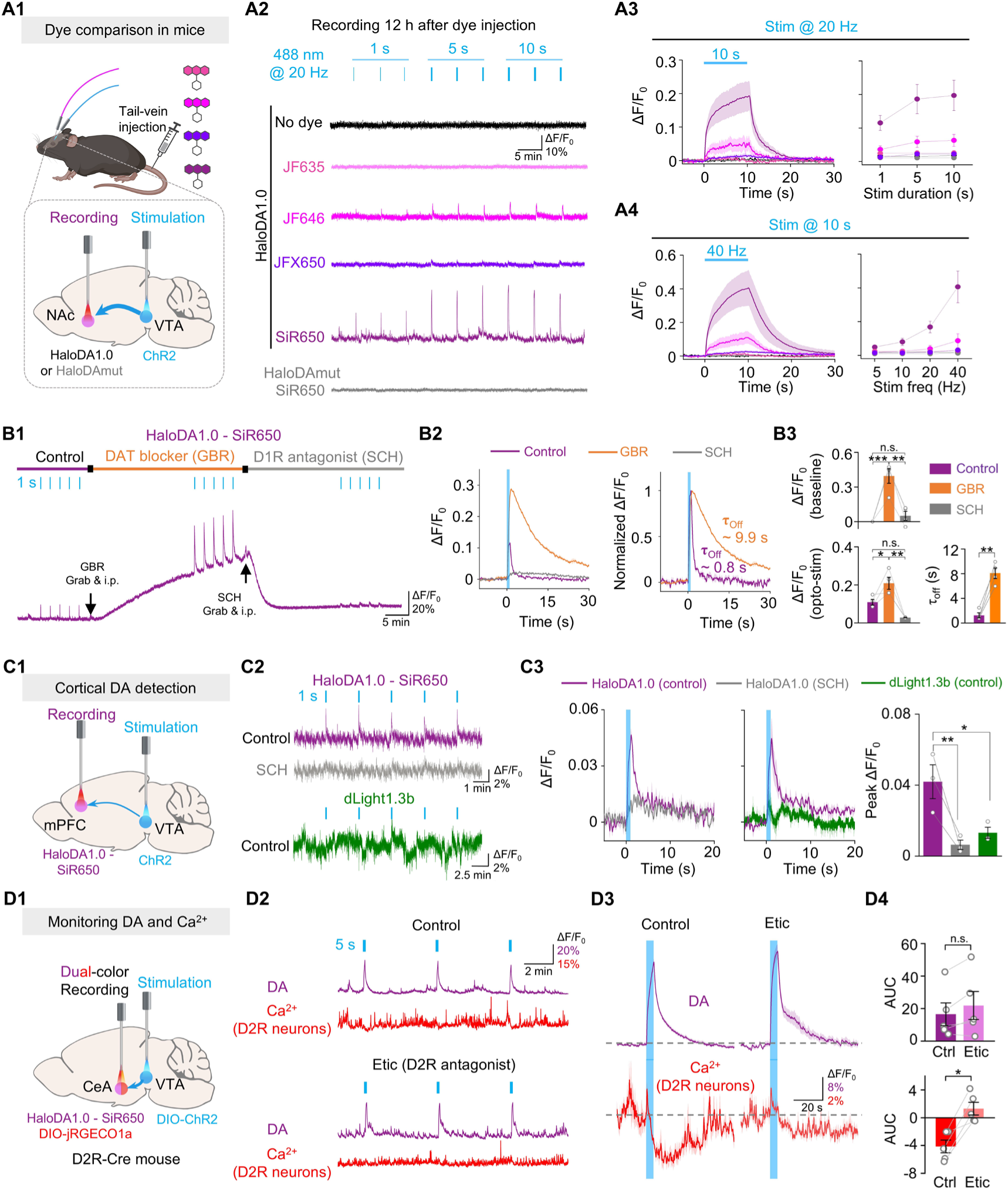
HaloDA1.0 can be used to detect endogenous DA release in freely moving mice. (**A1**) Schematic diagram depicting the strategy for using fiber photometry to record HaloDA1.0 or HaloDAmut labeled with various dyes (100 nmol injected via the tail vein) in the NAc upon optogenetic stimulation of VTA neurons. (**A2**) Representative traces of the change in HaloDA1.0 or HaloDAmut fluorescence during optogenetic stimulation in the indicated mice. (**A3** and **A4**) Average traces (left) and group summary (right) of the change in HaloDA1.0 or HaloDAmut fluorescence measured under the indicated stimulation duration (**A3**) and frequency (**A4**); n = 3–5 mice per group. (**B1**) Representative trace of the change in HaloDA1.0 fluorescence measured at baseline (control), after an intraperitoneal (i.p.) injection of 20 mg/kg GBR, and after an i.p. injection of 8 mg/kg SCH. The blue ticks indicate the optogenetic stimuli. (**B2**) Averaged trace (left) and normalized traces (right) of SiR650-labeled HaloDA1.0 measured in one mouse under the indicated conditions. The vertical blue shading indicates the optogenetic stimuli, and the off kinetics (τ_off_) were fitted with a single-exponential function. (**B3**) Group summary of baseline ΔF/F_0_ (top), peak ΔF/F_0_ (bottom left), and τ_off_ (bottom right) measured for SiR650-labeled HaloDA1.0 under the indicated conditions; n= 4 mice. (**C1**) Schematic illustration depicting the strategy for fiber photometry recording of HaloDA1.0 in the mPFC upon optogenetic stimulation of VTA neurons. (**C2**) Representative traces of the change in fluorescence of SiR650-labeled HaloDA1.0 and dLight1.3b under the indicated conditions. The blue ticks indicate the optogenetic stimuli applied at 20 Hz for 1 s. (**C3**) Average traces (5 trials averaged from one mouse on the left and 3 mice averaged on the middle) and group summary (right) of peak ΔF/F_0_ for SiR650-labeled HaloDA1.0 and dLight1.3b measured under the indicated conditions; n= 3 mice per group. The data for dLight1.3b were replotted from previously published results(*13*). (**D1**) Schematic diagram depicting the strategy for dual-color fiber photometry recording in the CeA with optogenetic stimulation of VTA neurons in a D2R-Cre mouse. (**D2** and **D3**) Representative traces (**D2**) and average traces (**D3**) of the DA and Ca^2+^ signals measured in D2R-expressing neurons in the same mouse under control conditions or following application of 2 mg/kg Etic. The blue ticks and vertical shading indicate the optogenetic stimuli applied at 20 Hz for 5 s. The traces in (**D3**) represent the average of 8 trials per condition. (**D4**) Group summary of the area under the curve (AUC, 0-30 s) of the DA and Ca^2+^ signals measured under the indicated conditions; n = 5 mice per group.

Next, we examined whether HaloDA1.0 can be used to monitor DA release *in vivo* in sparsely innervated brain regions such as the medial prefrontal cortex (mPFC)(*47*, *48*). We found that activation of neurons in the VTA caused transient increases in SiR650-labeled HaloDA1.0 in the mPFC, and these responses were blocked by SCH (Figs. 3C and S7B). In contrast, the genetically encoded green fluorescent DA sensor dLight1.3b(*15*) expressed in the mPFC did not show a measurable response to VTA stimulation (Fig. 3C), suggesting that unlike HaloDA1.0, dLight1.3b lacks the sensitivity needed to report DA release in the mPFC.

To test whether our far-red sensor is compatible for use in dual-color recordings during optogenetic stimulation, we expressed DIO-ChR2 in the VTA of D2R-Cre mice in order to specifically activate dopaminergic neurons, as D2R can serve as a general marker for these neurons in the VTA (*49*, *50*). In addition, we co-expressed HaloDA1.0 and the red fluorescent Ca^2+^ sensor DIO-jRGECO1a in the central nucleus of the amygdala (CeA)— which abundantly expresses DA receptors and receives dopaminergic projections from the VTA (*47*, *51*, *52*)—in order to examine how DA release affects neuronal activity in the CeA (Figs. 3D and S7C). We found that optogenetic stimuli triggered an increase in DA release together with a decrease in Ca^2+^ in D2R-positive neurons (Fig. 3D). Moreover, treatment with the D2R antagonist eticlopride blocked the change in Ca^2+^ without affecting DA release (Fig. 3D), indicating that DA may suppress the activity of D2R-positive neurons in the CeA by activating inhibitory D2R signaling.

### Simultaneously monitoring DA, ACh, and cAMP dynamics in the mouse NAc

In the striatum, both DA and ACh play essential roles in learning and motivation, regulating synaptic plasticity in part by binding to the excitatory D1 receptor and the inhibitory muscarinic acetylcholine M4 receptor (M4R), respectively, expressed on medium spiny neurons (D1-MSNs)(*53–57*). Although several pioneering studies examined the interaction between DA and ACh signaling(*8–10*), the effects of their concurrent regulation on intracellular cAMP signaling in D1-MSNs during behavior remain poorly understood. To address this important question, we virally co-expressed HaloDA1.0, rACh1h, and the green fluorescent cAMP sensor DIO-GFlamp2(*58*, *59*) in the NAc of D1R-Cre mice (Fig. 4A, B). We then labeled the DA sensor with SiR650 and used three-color fiber photometry to simultaneously monitor DA, ACh, and cAMP *in vivo* (Figs. 4 and S9).

**Fig. 4.**
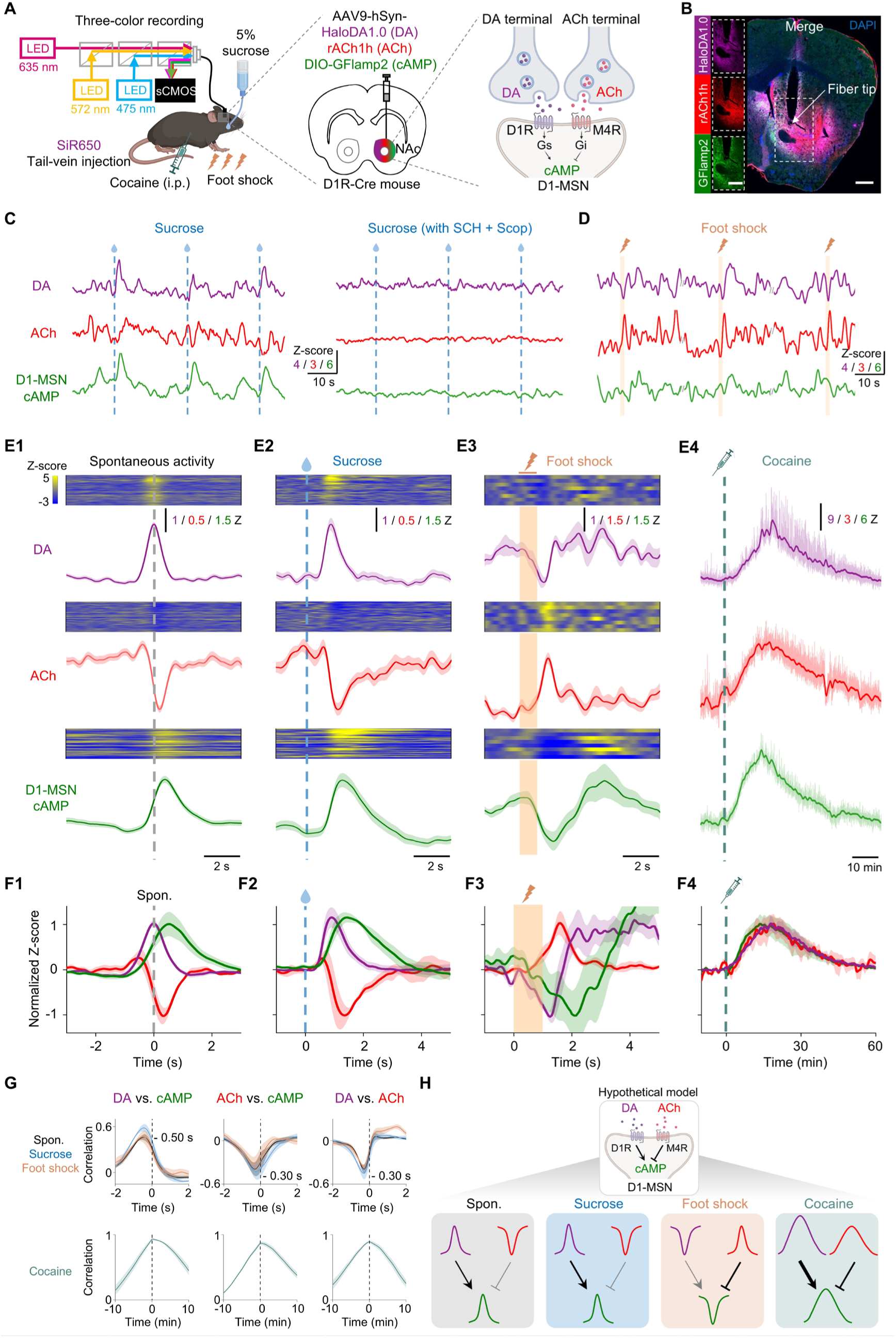
Simultaneous monitoring of DA, ACh, and cAMP dynamics *in vivo*. (**A**) (Left and middle) Schematic diagram depicting the strategy for three-color fiber photometry recording of DA, ACh, and D1-MSN cAMP signals in the NAc during 5% sucrose, foot shock (0.7 mA for 1 s), or following an i.p. injection of cocaine (20 mg/kg). (Right) Proposed model for the actions of DA and ACh in D1-MSNs. DA released from the dopaminergic termini binds Gs-coupled D1R to drive cAMP production. ACh released from cholinergic interneurons binds Gi-coupled M4R to reduce cAMP production. (**B**) Histological verification of HaloDA1.0, rACh1h, and DIO-GFlamp2 expression in the mouse NAc. The white arrow indicates the approximate location of the fiber tip. Images of the individual channels are shown on the left. Scale bars, 0.5 mm. (**C**) Example traces of all three sensor signals measured simultaneously in the NAc during three consecutive sucrose trials under control conditions (left) or after i.p. injection of 8 mg/kg SCH and 10 mg/kg Scop. (**D**) Example traces of all three sensor signals measured simultaneously in the NAc during three consecutive foot shock trials. (**E**) Representative time-aligned pseudocolor images and average traces of DA, ACh, and D1-MSN cAMP signals measured in a mouse during spontaneous activity (**E1**), sucrose (**E2**), foot shock (**E3**), and cocaine application (**E4**). The traces in **E1**, **E2**, and **E3** are shown as the mean ± s.e.m. In **E4**, the raw fluorescent response is indicated by the shaded area, and the bold lines indicate the response after low-pass filtering at 0.01 Hz. (**F**) Normalized fluorescence response of all three sensor signals measured during spontaneous activity (**F1**), sucrose (**F2**), foot shock (**F3**), and cocaine application (**F4**); n = 4 mice each. (**G**) Mean cross-correlation between the indicated pairs of sensor signals measured under the indicated conditions. The cross-correlations during spontaneous activity, sucrose, and foot shock application are shown in the top row, with the time lag indicated; n = 4 mice each. (**H**) Model illustrating the proposed effects of DA and ACh on D1-MSN cAMP levels. Elevated DA and reduced ACh increase cAMP production during spontaneous activity and in response to sucrose, while decreased DA and increased ACh reduce cAMP production during foot shock. In contrast, both DA and ACh increase in response to cocaine, exerting opposing effects on cAMP production.

All three signals showed spontaneous fluctuations under control conditions (i.e., in the absence of stimuli) (Fig. 4C). Centering on the peaks of the spontaneous DA fluctuations, we observed a corresponding increase in the cAMP signal and a phasic dip in the Ach signal (Fig. 4E1, F1). Interestingly, the peak in DA preceded the trough in the ACh signal, followed by the peak in cAMP, which is consistent with the requirement for DA to bind D1R in order to produce cAMP. During uncued sucrose rewards, we observed a pattern akin to the spontaneous signals (Fig. 4C, E2, F2); however, upon applying a 1-s foot shock, a distinct pattern emerged for all three signals (Fig. 4E3, F3). We then ruled out spectral crosstalk between the three sensors and confirmed the specificity of the signals by showing that SCH largely eliminated the DA and cAMP signals, while the M3R antagonist scopolamine selectively blocked the ACh signal (Fig. S10). Finally, a correlation analysis revealed a direct correlation between DA and cAMP (with a 500-ms lag) and inverse correlations both between ACh and cAMP and between DA and ACh (both with a 300-ms lag) (Fig. 4G), consistent with recent studies regarding the interaction between DA and ACh(*9*, *10*).

Combining their dynamics and receptor functions, we found that both the increase in DA and the decrease in ACh signals facilitate the production of cAMP during spontaneous activity and in response to sucrose (Fig. 4H). On the other hand, both the decrease in DA and the increase in ACh signals in response to aversive stimuli (in this case, foot shock) reduce cAMP production. We therefore examined the effect of the addictive drug cocaine on this regulatory mechanism. We found that a single i.p. injection of 20 mg/kg cocaine significantly increased all three signals, with the DA and cAMP signals being notably larger than the signals induced by sucrose (Figs. 4E4, F4 and S9). In addition, we found a strong direct correlation between all three pairs of signals (Fig. 4G), suggesting that cocaine can disrupt the normal interactions between these signaling processes (Fig. 4H). Together, these *in vivo* experiments provide a novel view of the dynamic interplay between DA, ACh, and cAMP in D1-MSNs under different conditions, highlighting the ability of using HaloDA1.0 to measure the sub-second dynamics and interplay between these neurochemicals *in vivo*.

## DISCUSSION

Here, we report the development, characterization, and application of HaloDA1.0, a far-red chemigenetic DA sensor with distinct spectral properties that make it compatible for use with existing sensors for monitoring other neuromodulators. By combining HaloDA1.0 with existing green and red fluorescent neuromodulator sensors, Ca^2+^ indicators, cAMP sensors, and optogenetic tools, we show that this DA sensor can be used for multi-color imaging in a variety of models. In cultured neurons, we simultaneously imaged the dynamics of three monoamines. In acute brain slices, we imaged the release—and we studied the regulation—of endogenous DA, ACh, and eCB upon electrical stimulation. Using zebrafish larvae, we imaged endogenous DA, ATP, and Ca^2+^ levels. Importantly, we also show that this sensor can detect DA release in mice, using dual-color *in vivo* imaging to measure changes in DA and intracellular Ca^2+^ in response to blue light–activated optogenetics. Finally, we simultaneously measured DA, ACh, and cAMP in the mouse NAc under basal conditions and during various behavioral stimuli, including sucrose, foot shock, and cocaine administration, revealing distinct patterns regulating these three signaling molecules.

Unlike genetically encoded DA sensors, HaloDA1.0 uses the chemical dye-cpHaloTag as its fluorescent module, in which the DA-dependent change in fluorescence relies on a shift in the equilibrium between dye’s L and Z forms. Despite using a different mechanism compared to conventional genetically encoded DA sensors, HaloDA1.0 has excellent sensitivity, good membrane trafficking, high specificity for DA, rapid kinetics, and minimal downstream coupling(*60*). Moreover, HaloDA1.0 can be used to monitor DA release in a wide range of brain regions, including the CeA and mPFC, making it superior to other sensors such as dLight1.3b, which lacks the necessary sensitivity to monitor DA release in the mPFC.

Although the cpHaloTag-based chemigenetic strategy by modulating L-Z equilibrium, has been used to develop both Ca^2+^ and voltage sensors(*25*, *26*), its *in vivo* applications have not yet been demonstrated. Identifying an appropriate dye for use *in vivo* is essential but challenging, requiring the right balance between its tunable properties and bioavailability. While some JF dyes, such as JF525 and JF669, demonstrate good blood-brain barrier permeability(*24*, *61*, *62*), and are compatible with a recent tryptophan quenching-based Ca^2+^ sensor(*63*), they were not suitable for labeling the HaloDA1.0 sensor. In addition, highly tunable rhodamine derivatives such as JF635 and JF646 have not yet been used *in vivo*. Here, we found that the dye SiR650 provided the best performance *in vivo,* presumably due to its high bioavailability. Future modifications to the dye’s structure and protein engineering are likely to further improve its labeling efficiency, achieving even better performance *in vivo*.

The ability to simultaneously measure DA, ACh, and cAMP in D1-MSNs within the NAc during various behaviors provides a highly comprehensive view of how neuromodulators and their downstream signals can integrate in order to modulate synaptic plasticity. Compared to previous studies that focused primarily on either the interaction between DA and ACh(*8–10*) or the interaction between DA and its downstream signals(*11*, *64*), our three-color recording system is more robust, yielding deeper insights than single-color and even dual-color recordings. Our findings suggest a potential synergistic modulation of D1- MSNs by DA and ACh under physiological conditions, and this delicate balance can be disrupted, for example by cocaine; this is consistent with previous studies showing that knocking out M4R in D1R-MSNs potentiates cocaine-induced hyperlocomotor activity(*65*).

Leveraging this chemigenetic strategy, we believe that in the future it will be possible to develop a wide range of far-red neuromodulator sensors based on other GPCRs. Given the more than 100 neurotransmitters and neuromodulators identified to date, this strategy will offer more options for researchers to simultaneously monitor multiple neurochemical signals. Moreover, by leveraging NIR dyes and protein engineering strategies(*23*, *24*, *66*), the sensors’ spectral range can be shifted even further into the NIR range, making them even more suitable for use in *in vivo* imaging. Ultimately, additional protein tags such as TMP-tag(*67*) and SNAP-tag(*68*) might be used to develop sensors that are orthogonal to existing cpHaloTag sensors, providing the ability to simultaneously image a multitude of neuromodulators.

In summary, our far-red chemigenetic DA sensor, which is suitable for both *in vitro* and *in vivo* applications, can be used to simultaneously measure multiple neurochemical signals in real time. This robust new tool can therefore be used to significantly increase our understanding of the regulatory mechanisms and specific roles of the dopaminergic system under both physiological and pathological conditions.

## METHODS

### Molecular biology

Plasmids were generated using the Gibson assembly method. Primers for PCR amplification of DNA fragments were synthesized (Ruibio Biotech) with 30-base pair overlap. The cDNA encoding D1R was cloned from the human GPCR cDNA library (hORFeome database 8.1), and the cDNA encoding cpHaloTag was synthesized (Shanghai Generay Biotech) based on the reported sequence(*25*). All constructs were verified using Sanger sequencing (Ruibio Biotech and Tsingke Biotech).

For screening and characterization in HEK293T cells, cDNAs encoding the candidate sensors were cloned into a modified pDisplay vector (Invitrogen) containing an upstream IgK leader sequence, followed by an IRES and membrane-anchored EGFP-CAAX for calibration. Site-directed mutagenesis was performed using primers with randomized NNS codons (32 codons in total, encoding all 20 possible amino acids). To measure the spectra, a stable cell line was generated by cloning the HaloDA1.0 gene into the pPacific vector, which contains a 3′ terminal repeat, IRES, the puromycin gene, and a 5’ terminal repeat. For the luciferase complementation assay, the D1R/HaloDA1.0-SmBit construct was created by replacing the β2AR gene in β2AR-SmBit with D1R or HaloDA1.0, and miniGs- LgBit was generously provided by N. A. Lambert (Augusta University). For the Tango assay, D1R-Tango was cloned from the PRESTO-Tango GPCR Kit (Addgene kit no. 1000000068), and HaloDA1.0-Tango was generated by replacing D1R in D1R-Tango with HaloDA1.0. For characterization in cultured neurons, acute brain slices, and *in vivo* mouse experiments, the HaloDA1.0 and HaloDAmut sensors were cloned into the pAAV vector under the control of the human *Synapsin* promoter and used for AAV packaging. For zebrafish imaging, the HaloDA1.0 and HaloDAmut sensors were cloned into elavl3:Tet^off^ vectors, followed by P2A- EGFP or independent EGFP expression under the control of the zebrafish *myl7* promoter.

### Preparation and fluorescence imaging of cultured cells

#### Cell culture and transfection

The HEK293T cell line was purchased from ATCC (CRL-3216) and cultured in high-glucose Dulbecco’s Modified Eagle’s Medium (Gibco) supplemented with 10% (v/v) fetal bovine serum (CellMax) and 1% penicillin-streptomycin (Gibco) at 37°C in humidified air containing 5% CO_2_. For screening and characterizing the sensors, the cells were plated on 96-well plates and grown to 70% confluence before transfection with a mixture containing 0.3 μg DNA and 0.9 μg 40-kDa polyethylenimine (PEI) for 6-8 h. For kinetics measurements, cells were plated on 12-mm glass coverslips in 24-well plates and transfected with a mixture containing 1 μg DNA and 3 μg PEI for 6-8 h. Fluorescence imaging was conducted 24-36 h after transfection.

Rat primary cortical neurons were prepared from postnatal day 0 (P0) Sprague-Dawley rat pups (Beijing Vital River Laboratory) and dissociated using 0.25% trypsin-EDTA (Gibco). The neurons were plated on 12-mm glass coverslips coated with poly-D-lysine (Sigma-Aldrich) in 24-well plates and cultured with Neurobasal medium (Gibco) supplemented with 2% B-27 (Gibco), 1% GlutaMAX (Gibco), and 1% penicillin-streptomycin (Gibco) at 37 °C in humidified air containing 5% CO_2_. Every 3 days, 50% of the media was replaced with fresh media. At 3 days in culture (DIV3), cytosine β-D-arabinofuranoside (Sigma) was added to the cortical cultures to a final concentration of 1 μM. For characterization in cultured neurons, cortical cultures were transduced with adeno-associated virus (AAV) expressing HaloDA1.0 (full titer, 1 μl per well) at DIV6 and imaged at DIV15-20. For three-color neuron imaging, AAVs expressing HaloDA1.0, r5-HT1.0, and NE2m (full titer, 1 μl per well for each virus) were sequentially added to the cortical cultures at DIV6, DIV9, and DIV12, respectively, to minimize expression competition, and imaging was performed at DIV20-23.

#### Imaging of HEK293T cells

Before imaging, HEK293T cells expressing HaloDA1.0—or variants thereof—were pre-treated with 0.5-1 μM dye for 1 h, followed by washing with fresh culture medium for an additional 2 h. The culture medium was then replaced with Tyrode’s solution consisting of (in mM): 150 NaCl, 4 KCl, 2 MgCl_2_, 2 CaCl_2_, 10 HEPES, and 10 glucose (pH adjusted to 7.35-7.45 with NaOH). HEK293T cells plated on 96-well CellCarrier Ultra plates (PerkinElmer) were imaged using the Operetta CLS high-content analysis system (PerkinElmer) equipped with a 20×, numerical aperture (NA 1.0) water-immersion objective and an sCMOS camera to record fluorescence. A 460-490-nm LED and 500-550-nm emission filter were used to image green fluorescence (e.g., EGFP); a 530-560-nm LED and 570-620-nm emission filter were used to image yellow fluorescence (e.g., JF525 and JF526); a 530-560-nm LED and 570-650 nm emission filter were used to image red fluorescence (e.g., JF585); a 615-645-nm LED and 655-760-nm emission filter were used to image far-red fluorescence (e.g., JF635, JF646, JFX650, and SiR650); and a 650-675- nm LED and 685-760-nm emission filter were used to image near-infrared fluorescence (e.g., SiR700).

During imaging, the following compounds were applied via bath application at the indicated concentrations: DA (Sigma-Aldrich), SCH-23390 (MedChemExpress), eticlopride (Tocris), SKF-81297 (Tocris), quinpirole (Tocris), serotonin (Tocris), histamine (Tocris), octopamine (Tocris), tyramine (Sigma-Aldrich), ACh (Solarbio), γ-aminobutyric acid (Tocris), glutamate (Sigma-Aldrich), levodopa (Abcam), and NE (Tocris). The fluorescence signals produced by the HaloDA1.0 sensors were calibrated using EGFP, and the change was in fluorescence (ΔF/F_0_) was calculated using for formula (F - F_0_)/F_0_, where F_0_ is the baseline fluorescence.

#### Imaging of cultured neurons

Before imaging, cultured neurons expressing HaloDA1.0 were pre-treated for 1 h with 1 μM JF635 or JF646, or with 200 nM SiR650 or JFX650 to minimize non-specific labeling. The dyes were then removed by washing the neurons with culture medium for an additional 2-3 h, and Tyrode’s solution was used for imaging. The neurons, plated on 12-mm glass coverslips, were bathed in a custom-made chamber for imaging using an inverted A1R Si+ laser scanning confocal microscope (Nikon) equipped with a 20× (NA: 0.75) objective and a 40× (NA: 1.35) oil-immersion objective. A 488-nm laser and 525/50-nm emission filter were used to image green fluorescence (e.g., NE2m); a 561-nm laser and 595/50-nm emission filter were used to image red fluorescence (e.g., r5-HT1.0); and a 640-nm laser and 700/75-nm emission filter were used to image far-red fluorescence (e.g., HaloDA1.0 labeled with JF635, JF646, SiR650, or JFX650). For single-color imaging, images were acquired with a frame interval of 5 s. For three-color imaging, the fluorescence signals from the green, red and far-red sensors were acquired sequentially, with a period interval of 5 s. The change in fluorescence (ΔF/F_0_) was calculated using the formula (F - F_0_)/F_0_.

#### Kinetics measurements

HEK293T cells expressing HaloDA1.0 were plated on 12-mm glass coverslips, labeled with JF646 or SiR650, and imaged using an A1R confocal microscope (Nikon) equipped with a 40× (NA: 1.35) oil-immersion objective. A glass pipette was positioned approximately 10- 20 μm from the sensor-expressing cells, and fluorescence signals were recorded using the confocal high-speed line scanning mode at a scanning frequency of 1,024 Hz. To measure τ_on_, 100 μM DA was puffed onto the cells from the pipette, and the resulting increase in fluorescence was fitted with a single-exponential function. To measure τ_off_, 100 μM SCH- 23390 was puffed onto cells bathed in 1 μM DA, and the resulting decrease in fluorescence was fitted with a single-exponential function.

### Tango assay

HTLA cells stably expressing a tTA-dependent luciferase reporter and a β-arrestin2-TEV gene were a gift from B.L. Roth (University of North Carolina Medical School). The cells were initially plated in 6-well plates and transfected with either HaloDA1.0-Tango or D1R- Tango; 24 h after transfection, the cells were transferred to 96-well plates and incubated with varying concentrations of DA (ranging from 0.01 nM to 100 µM). In addition, 1 µM JF646 was applied to half of the wells. The cells were then cultured for 12 h to allow expression of tTA-dependent luciferase. Bright-Glo reagent (Fluc Luciferase Assay System, Promega) was added to a final concentration of 5 µM, and luminescence was measured 10 min later using a VICTOR X5 multi-label plate reader (PerkinElmer).

### Mini G protein luciferase complementation assay

HEK293T cells were first plated in 6-well plates and co-transfected with a pcDNA3.1 vector expressing either HaloDA1.0-SmBit or D1R-SmBit (or empty vector) together with miniGs- LgBit; 24 h after transfection, the cells were dissociated and mixed with Nano-Glo Luciferase Assay Reagent (Promega) diluted 1,000-fold to a final concentration of 5 µM. The cell suspension was then distributed into 96-well plates and treated with various concentrations of DA. Following a 10-min incubation in the dark, luminescence was measured using a VICTOR X5 multi-label plate reader (PerkinElmer).

### Spectra measurements

#### One-photon spectral characterization

The one-photon spectra were measured using a Safire 2 microplate reader (Tecan). HEK293T cells stably expressing HaloDA1.0 were plated in 6-well plates and labeled with dye after 24 h. The cells were then harvested and transferred to black 384-well plates. The fluorescence values measured in unlabeled cells were subtracted as background. Both the excitation and emission spectra were measured in the presence of saline or 100 μM DA at 5-nm increments. Below are the wavelength settings for each dye-labeled sample:

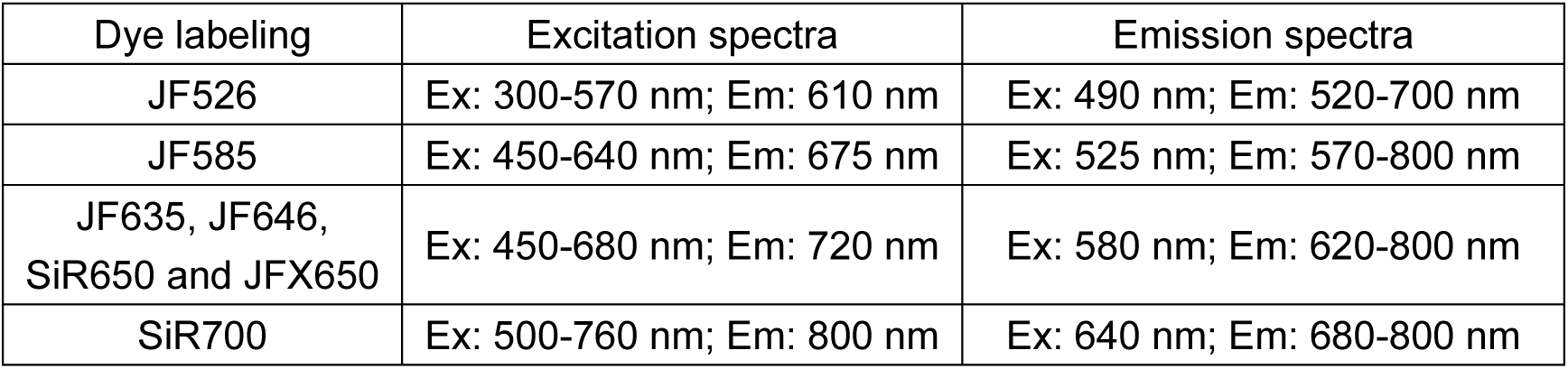

#### Two-photon spectral characterization

HEK293T cells expressing HaloDA1.0 were plated on 12-mm glass coverslips and labeled with JF646 or SiR650. Two-photon excitation spectra were measured at 10-nm increments ranging from 870 nm to 1300 nm using an Olympus FVMPE-RS microscope equipped with a tunable Spectra-Physics InSight X3 laser. The far-red signals were collected with a 660- 750-nm emission filter and a 760-nm dichroic mirror positioned between the lasers and photomultiplier tubes (PMTs). The recorded signals were calibrated according to the output power of the tunable two-photon laser at each wavelength.

### Synthesis of chemical dyes

#### Synthesis of SiR650-HTL

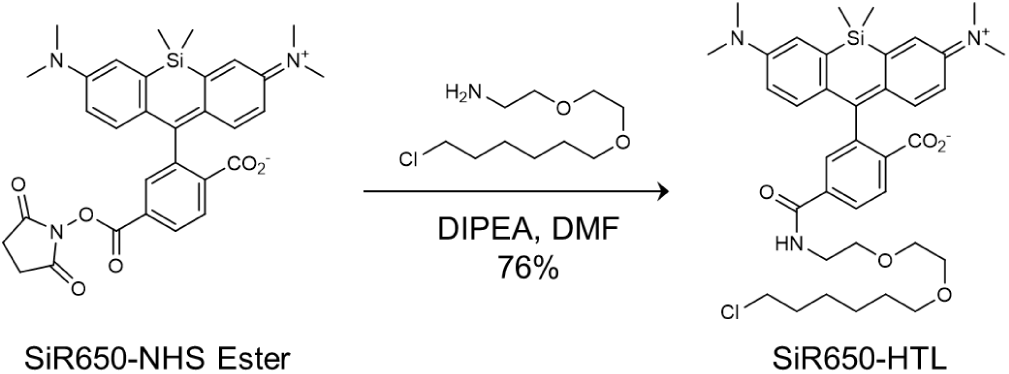

SiR-NHS Ester (23 mg, 40 μmol, 1.0 eq., obtained from CONFLUORE) and HaloTag(O2)amine (13 mg, 60 μmol, 1.5 eq.) were dissolved in 2 ml anhydrous DMF. DIPEA (13 μl, 80 μmol, 2.0 eq.) was then added, and the mixture was stirred at room temperature overnight. Purification of the mixture by reverse phase-HPLC (eluent, a 30- min linear gradient, from 20% to 95% solvent B; flow rate, 5.0 mL/min; detection wavelength, 650 nm; eluent A (ddH_2_O containing 0.1% TFA (v/v)) and eluent B (CH_3_CN)) provided SiR650-HTL (21 mg, 76% yield) as a blue solid.

^1^H NMR (DMSO-*d*_6_, 400 MHz) δ 8.78 (t, *J* = 5.5 Hz, 1H), 8.08 (dd, *J* = 8.0, 1.3 Hz, 1H), 8.02 (dd, *J* = 8.0, 0.4 Hz, 1H), 7.69 – 7.65 (m, 1H), 7.03 (d, *J* = 2.4 Hz, 2H), 6.65 (dd, *J* = 9.0, 2.6 Hz, 2H), 6.61 (d, *J* = 8.9 Hz, 2H), 3.57 (t, *J* = 6.7 Hz, 2H), 3.53 – 3.46 (m, 4H), 3.46 – 3.40 (m, 2H), 3.40 – 3.34 (m, 2H), 3.30 (t, *J* = 6.5 Hz, 2H), 2.94 (s, 12H), 1.70 – 1.60 (m, 2H), 1.46 – 1.36 (m, 2H), 1.36 – 1.19 (m, 4H), 0.65 (s, 3H), 0.53 (s, 3H). Analytical HPLC, > 99% purity (4.6 mm × 150 mm 5 μm C18 column; 2 μl injection; 5-100% CH_3_CN/H_2_O, linear-gradient, with constant 0.1% v/v TFA additive; 6 min run; 0.6 ml/min flow; ESI; positive ion mode; detection at 650 nm). HRMS (ESI) calcd for C_37_H_49_ClN_3_O_5_Si [M+H]^+^ 678.3130, found 678.3133.

#### Synthesis of JF646-HTL

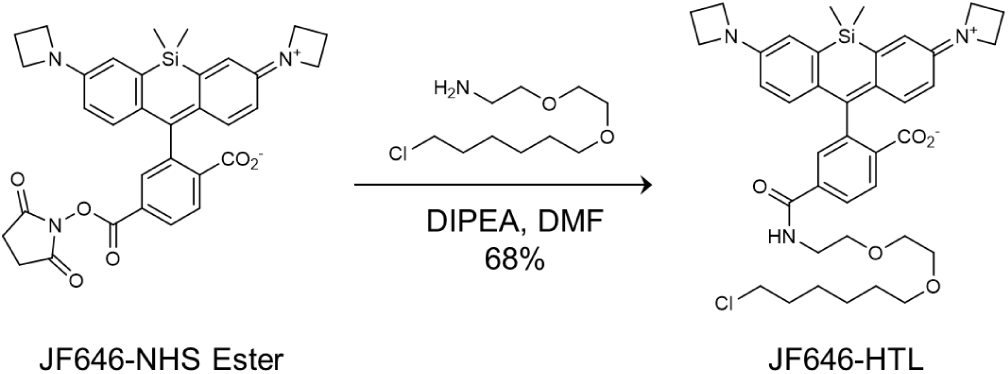

JF646-NHS Ester (24 mg, 40 μmol, 1.0 eq., obtained from AAT Bioquest) and HaloTag(O2)amine (13 mg, 60 μmol, 1.5 eq.) were dissolved in 2 ml anhydrous DMF. DIPEA (13 μl, 80 μmol, 2.0 eq.) was then added, and the mixture was stirred at room temperature overnight. Purification of the mixture by reverse phase-HPLC (eluent, a 30- min linear gradient, from 20% to 95% solvent B; flow rate, 5.0 ml/min; detection wavelength, 650 nm; eluent A (ddH_2_O containing 0.1% TFA (v/v)) and eluent B (CH_3_CN)) provided JF646-HTL (19 mg, 68% yield) as a blue solid.

^1^H NMR (CDCl_3_, 400 MHz) δ 8.00 (dd, *J* = 8.0, 0.7 Hz, 1H), 7.92 (dd, *J* = 8.0, 1.3 Hz, 1H), 7.70 (dd, *J* = 1.2, 0.7 Hz, 1H), 6.75 (d, *J* = 8.7 Hz, 2H), 6.73 – 6.67 (m, 1H), 6.65 (d, *J* = 2.6 Hz, 2H), 6.26 (dd, *J* = 8.8, 2.7 Hz, 2H), 3.89 (t, *J* = 7.3 Hz, 8H), 3.67 – 3.60 (m, 6H), 3.56 – 3.53 (m, 2H), 3.50 (t, *J* = 6.5 Hz, 2H), 3.39 (t, *J* = 6.6 Hz, 2H), 2.39 – 2.30 (m, 4H), 1.78 – 1.68 (m, 2H), 1.56 – 1.47 (m, 2H), 1.44 – 1.35 (m, 2H), 1.35 – 1.25 (m, 2H), 0.63 (s, 3H), 0.56 (s, 3H). Analytical HPLC, > 99% purity (4.6 mm × 150 mm 5 μm C18 column; 2 μl injection; 5-100% CH_3_CN/H_2_O, linear-gradient, with constant 0.1% v/v TFA additive; 6 min run; 0.6 mL/min flow; ESI; positive ion mode; detection at 650 nm). HRMS (ESI) calcd for C_39_H_49_ClN_3_O_5_Si [M+H]^+^ 702.3125, found 702.3140.

### Mice and viruses

Wild-type C57BL/6J mice of both sexes (6-10 weeks of age) were obtained from Beijing Vital River Laboratory. D2R-Cre mice were kindly provided by M. Luo at the Chinese Institute for Brain Research, Beijing, and D1R-Cre mice were kindly provided by Y. Rao at Peking University. All animal protocols were approved by the Animal Care and Use Committee at Peking University. All animals were housed under a 12-h/12-h light/dark cycle at an ambient temperature of 25°C and were provided food and water ad libitum.

For dye injection in mice, unless otherwise noted, the following formulation was used: 20 μl of 5 mM SiR650 or other far-red dye (in DMF, equivalent to 100 nmol) was mixed with 20 μl Pluronic F-127 (20% w/v in DMSO, AAT Bioquest) and 100 μl PBS and injected via the tail vein the day before recording or imaging.

The following viruses were packaged at Vigene Biosciences: AAV9-hSyn-HaloDA1.0 (7.73×10^13^ viral genomes (vg)/ml), AAV9-hsyn-hChR2(H134R)-mCherry (2.53×10^13^ vg/ml), AAV9-EF1α-DIO-hChR2(H134R)-EYFP (9.12×10^13^ vg/ml), AAV9-hSyn-NE2m (1.39×10^13^ vg/ml), and AAV9-hSyn-r5-HT1.0 (1.06×10^13^ vg/ml). AAV-hsyn-haloDA1.0mut (5.38×10^12^ vg/ml) was packaged at BrainVTA. In addition, the following two viruses were co-packaged at BrainVTA with mixed plasmids (1:1:1 ratio) to reduce mutual suppression: AAV9-hSyn-HaloDA1.0 / AAV9-hsyn-rACh1h / AAV9-hsyn-DIO-GFlamp2 (5.54×10^12^ vg/ml) and AAV9- hSyn-HaloDA1.0 / AAV9-hsyn-rACh1h / AAV9- hsyn-eCB2.0 (5.83×10^12^ vg/ml). AAV9- EF1α-DIO-NES-jRGECO1a (5.76×10^12^ vg/ml) was packaged at Brain Case.

### Fluorescence imaging of acute brain slices

#### Preparation of brain slices

Adult male C57BL/6J mice (8-10 weeks old) were anesthetized via intraperitoneal injection of 2,2,2-tribromoethanol (Avertin, 500 mg/kg). A stereotaxic injection of AAV9-hSyn-HaloDA1.0 (300 nl) or a co-packaged virus containing AAV9-hSyn-HaloDA1.0, AAV9- hSyn-rACh1h, and AAV9-hSyn-eCB2.0 (400 nl total volume) was delivered into the nucleus accumbens (NAc) core at a rate of 50 nl/min. The injection coordinates were: AP +1.4 mm relative to Bregma, ML ±1.2 mm relative to Bregma, and DV-4.3 mm from the dura. After 2-4 weeks, the mice were deeply anesthetized, followed by transcardiac perfusion with cold slicing buffer consisting of (in mM): 110 choline chloride, 2.5 KCl, 1.25 NaH_2_PO_4_, 25 NaHCO_3_, 7 MgCl_2_, 25 glucose, and 0.5 CaCl_2_. The brain was quickly extracted, placed in cold, oxygenated slicing buffer, and sectioned into 300-μm coronal slices using a VT1200 vibratome (Leica).

For imaging of JF646-labeled slices, the brain slices were first incubated in oxygenated ACSF containing 1 µM JF646 at room temperature for 60 min. The ACSF contained (in mM): 125 NaCl, 2.5 KCl, 1 NaH_2_PO_4_, 25 NaHCO_3_, 1.3 MgCl_2_, 25 glucose, and 2 CaCl_2_. After incubation, the slices were transferred to fresh oxygenated ACSF and allowed to sit for at least 60 min to remove any non-specific dye binding. For imaging of SiR650-labeled slices, 100 nmol of SiR650 was injected into the tail vein; 12 h after injection, acute brain slices were prepared as described above and incubated in oxygenated ACSF for at least 60 min at room temperature before imaging.

#### Single-color imaging of acute brain slices

Confocal imaging was conducted using a Zeiss LSM-710 confocal microscope equipped with a N-Achroplan 20x (NA: 0.5) water-immersion objective, a HeNe633 laser, a HeNe543 laser, and an Argon laser. The microscope was controlled using ZEN2012, 11.0.4.190 software (Zeiss). Slices were mounted in a custom-made imaging chamber with continuous ACSF perfusion at 2 ml/min.

HaloDA1.0 labeled with JF646 was excited using a 633-nm laser, and fluorescence emission captured at 638-747 nm. Images were acquired at a size of 256 × 256 pixels and a frame rate of 5 Hz. Electrical stimuli were applied using a Grass S48 stimulator (Grass Instruments). A bipolar electrode (WE30031.0A3, MicroProbes) was placed near the NAc core under fluorescence guidance, and stimuli were applied at a voltage of 4-7 V and a pulse duration of 1 ms. Synchronization of imaging and stimulation was facilitated using an Arduino board (Uno) with custom scripts controlling the process. To calculate ΔF/F_0_, baseline fluorescence was defined as the average fluorescence signal obtained for 10 s before stimulation.

For kinetics measurements, a zoomed-in region (64 x 64 pixels) was scanned at a frame rate of 13.5 Hz. A single 1-ms pulse was delivered, and resulting increase and subsequent decrease in fluorescence were fitted with single-exponential functions.

#### Three-color imaging of acute brain slices

Three-color imaging of acute brain slices was performed using a Zeiss LSM-710 confocal microscope, with the signals from three sensor captured in two sequential scans in order to minimize spectral interference. First, we simultaneously imaged HaloDA1.0 and eCB2.0; we then performed a separate scan to image rACh1h. HaloDA1.0 was excited at 633 nm, and the emitted fluorescence was captured at 645-700 nm; eCB2.0 was excited at 488 nm, and the emitted fluorescence was captured at 509-558 nm; finally, rACh1h was excited at 543 nm, and the emitted fluorescence was captured at 580-625 nm. Images were acquired at 256 × 256 pixels at a frequency of 4 Hz. The change in fluorescence was calculated as described above, with the baseline calculated using as the average fluorescence signal measured for 0-10 s before stimulation.

Field stimuli (1-ms duration) were applied using parallel platinum electrodes (1 cm apart), with voltage ranging from 40-80 V. During imaging, the following compounds were added to the imaging chamber at a rate of 2 ml/min: SCH-23390 (MedChemExpress), scopolamine (MedChemExpress), AM251 (Cayman), GBR12909 (MedChemExpress), and donepezil (MedChemExpress).

### Fluorescent imaging of zebrafish larvae

For these experiments, we used 4-6 days post-fertilization (dpf) zebrafish larvae. Before imaging, the larvae were immersed in dye (3.3 μM) for 1 h, then transferred to plain water for 2 h to remove the dye from the larvae’s surface. Zebrafish embryos and larvae were maintained at 28°C on a 14-h light and 10-h dark cycle. All procedures were approved by the Institute of Neuroscience, Chinese Academy of Sciences.

#### Comparison of various dyes in zebrafish

For single-channel imaging, the elavl3:Tet^off^-HaloDA1.0-P2A-EGFP or elavl3:Tet^off^- HaloDAmut-P2A-EGFP plasmid (25 ng/μl) mixed with Tol2 transposase mRNA (25 ng/μl) was injected into fertilized embryos on a Nacre (*mitfa^w2/w2^*) background at the one-cell stage in order to generate chimeric transgenic fish. Positive fish were selected based on EGFP expression. After being labeled with dye, the zebrafish were embedded in 2% agarose gel and imaged using an FN1 confocal microscope (Nikon) equipped with a 16x (NA: 0.8) water-immersion objective. HaloDA1.0 and HaloDAmut were excited using a 640- nm laser, and fluorescence emissions were captured at 650-750 nm. Time-lapse images were acquired at 512 x 512 pixels at ∼1.06 s per frame. PBS, either with or without 100 μM DA, was locally puffed on the larvae using a micropipette with a tip diameter of 1-2 μm, targeting the optic tectum region. The change in fluorescence was calculated as described above, with the baseline calculated using the average fluorescence signal measured for 10-50 s before puff application.

#### Three-color imaging in zebrafish

For three-color imaging, the HaloDA1.0 plasmid, which co-expresses the cardiac green fluorescent marker myl7-EGFP to facilitate the selection of positive fish, was injected into double transgenic embryos in order to generate chimeric triple transgenic larval zebrafish. Specifically, the elavl3:Tet^off^-HaloDA1.0;myl7-EGFP plasmid (25 ng/μl) mixed with Tol2 transposase mRNA (25 ng/μl) were injected into Tg(gfap:Tet^off^- ATP1.0);Tg(elavl3:jRGECO1a) embryos. An FV3000 confocal microscope (Olympus) equipped with a 20x (NA: 1.0) water-immersion objective was used for imaging. HaloDA1.0 was excited at 640 nm, and the emitted fluorescence was captured at 650-750 nm; jRGECO1a was excited at 561 nm, and the emitted fluorescence was captured at 570-620 nm; finally, ATP1.0 was excited at 488 nm, and the emitted fluorescence was captured 500- 540 nm. Time-lapse imaging was performed using the sequential line-scanning mode in order to obtain three sensor images (512 x 512 pixels) at a frame rate of 0.5 Hz.

Electric shock was generated using an ISO-Flex stimulus isolator (A.M.P.I), controlled by a programmable Arduino board (Uno), and applied using silver-plated tweezers placed parallel to the fish. Each stimulus was applied at 40 V/cm, with a duration of 1 s and an interval of 180 s. The change in fluorescence was calculated as described above, with the baseline calculated using the average fluorescence signal measured for 0-30 s before electrical shock. The cross-correlation between each pair of signals (DA, Ca^2+^, and/or ATP) in Fig. S6D was calculated using the *xcorr* function in MATLAB. Similar cross-correlation analysis was also applied to the three-color fiber photometry data (Fig. 4G).

In the PTZ imaging experiment, the baseline responses were recorded for 5 min, followed by the addition of PTZ to a final concentration of 10 mM, and imaging was continued for 0.5-1 h. To identify the peak in Fig. S6E, the Ca^2+^ peak was selected using the MATLAB *findpeaks* function with a minimum peak prominence set to one-tenth of the maximum Ca^2+^ response for each zebrafish. For adjacent peaks with an interval <70 s, only the highest peak was selected. The DA, Ca^2+^, and ATP transients were aligned to the Ca^2+^ peak. Peaks were further selected only if the peak amplitude of ATP and DA was exceeded one-tenth of the maximum response for each zebrafish.

### *In vivo* fiber photometry recording with optogenetic stimulation in mice

#### Optogenetic recording in the NAc and mPFC

Adult male C57BL/6J mice (8-10 weeks old) were anesthetized, and AAV9-hsyn-HaloDA1.0 or AAV9-hsyn-HaloDAmut (300 nl) was injected into the NAc (AP: +1.4 mm relative to Bregma, ML: ±1.2 mm relative to Bregma, and DV:-4.0 mm from the dura) or mPFC (AP: +1.98 mm relative to Bregma, ML: ±0.3 mm relative to Bregma, and DV:-1.8 mm from the dura). Virus expressing AAV9-hsyn-hChR2(H134R)-mCherry (500 nl) was injected into the ipsilateral VTA (AP:-2.9 mm relative to Bregma, ML: ±0.6 mm relative to Bregma, and DV:-4.1 mm from the dura). An optical fiber (200-μm diameter, 0.37 NA; Inper) was implanted 0.1 mm above the virus injection site in the NAc or mPFC, and another optical fiber was implanted 0.2 mm above the virus injection site in the VTA.

At 2-3 weeks after virus injection, the mice were injected with various far-red dyes; 12 h after injection, photometry recording with optogenetic stimulation was performed. The sensor signals were recorded using a customized photometry system (Thinker Tech) equipped with a 640/20-nm bandpass-filtered (Model ZET640/20x; Chroma) LED light (Cree LED) for excitation; a multi-bandpass-filtered (Model ZET405/470/555/640m; Chroma) PMT (Model H10721-210; Hamamatsu) was used to collect the signal, and an amplifier (Model C7319; Hamamatsu) was used to convert the current output from the PMT to a voltage signal. The voltage signal was passed through a low-pass filter and then digitized using an acquisition card (National Instruments). The excitation light power at the tip of the optical fiber was 80 μW and was delivered at 20 Hz with a 10-ms pulse duration.

An external 488-nm laser (LL-Laser) was used for optogenetic stimulation and was controlled by the photometry system to allow for staggered stimulation and signal recording. The stimulation light power at the tip of the fiber was 20 mW, and 10-ms pulses were applied. Three stimulation patterns were used: stimuli were applied for 1 s, 5 s, or 10 s at 20 Hz; stimuli were applied at 5 Hz, 10 Hz, 20 Hz, or 40 Hz for 10 s; and stimuli were applied for fixed duration (1 s) and frequency (20 Hz). Where indicated, the mice received an intraperitoneal injection of SCH-23390 (8 mg/kg) or GBR12909 (20 mg/kg) in a total volume of 300-400 μl. ΔF/F_0_ was calculated as described above, with the baseline calculated as the average fluorescence signal measured for 15-30 s before optogenetic stimulation.

#### Dual-color optogenetic recording in the CeA

Adult male and female D2R-Cre mice (8-12 weeks old) were used for this experiment. A 2:1 mixture of AAV9-hSyn-HaloDA1.0 and AAV9-EF1α-DIO-NES-jRGECO1a (400 nl total volume) was injected into the CeA (AP:-1 mm relative to Bregma, ML: ±2.5 mm relative to Bregma, and DV:-4.3 mm from the dura). AAV9-EF1α-DIO-hChR2(H134R)-EYFP (400 nl) was also injected into the ipsilateral VTA (AP:-2.9 mm relative to Bregma, ML: ±0.6 mm relative to Bregma, and DV:-4.1 mm from the dura). Two optical fibers (200-μm diameter, 0.37 NA; Inper) were implanted 0.1 mm above the virus injection site in the CeA and 0.2 mm above the virus injection site in the VTA.

Three weeks after virus injection, a customized three-color photometry system (Thinker Tech) was used for photometry recording as described in the following section. The system was equipped with three LEDs, but only two LEDs were used in this experiment to excite the red fluorescent jRGECO1a sensor at 40 μW and the far-red HaloDA1.0 sensor at 80 μW. The excitation lights were delivered sequentially at 20 Hz with a 10-ms pulse duration for each. An external 473-nm laser (LL-Laser) was used for optogenetic stimulation and was controlled by the photometry system to allow for staggered stimulation and signal recording. The stimulation light power at the tip of the fiber was 20 mW, with a 10-ms duration for each pulse. The day before recording, the mice received an injection of SiR650 via the tail vein. Where indicated, the mice also received an intraperitoneal injection of eticlopride (2 mg/kg) at a total volume of 350 μl. ΔF/F_0_ was calculated as described above, and, the baseline was calculated as the average fluorescence signal measured for 15-30 s before optogenetic stimulation. The area under the curve (AUC) in Fig. 3D was calculated during the 0-30 s period after optogenetic stimulation.

### *In vivo* three-color recording in the NAc

Adult male and female D1R-Cre mice (10-14 weeks old) were used for this experiment. A co-packaged AAV mixture containing AAV9-hSyn-HaloDA1.0, AAV9-hsyn-rACh1h, and AAV9-hsyn-DIO-GFlamp2 (600 nl total volume) was unilaterally injected into the NAc (AP: +1.4 mm relative to Bregma, ML: ±1.2 mm relative to Bregma, and DV:-4.0 mm from the dura), and an optical fiber (200-μm diameter, 0.37 NA; Inper) was implanted 0.1 mm above the virus injection site.

Photometry recording was performed 2-3 weeks after virus injection using a customized three-color photometry system (Thinker Tech). A 470/10-nm (model 65144; Edmund optics) filtered LED at 40 μW was used to excite the green fluorescent sensors; a 555/20-nm (model ET555/20x; Chroma) filtered LED at 40 μW was used to excite the red fluorescent sensors; and a 640/20-nm (model ZET640/20x; Chroma) filtered LED at 40 μW was used to excite the far-red fluorescent sensors. The three excitation lights were delivered sequentially at 20-Hz with a 10-ms pulse duration for each, and fluorescence was collected using an sCMOS (Tucsen) and filtered with a three-bandpass filter (model ZET405/470/555/640m; Chroma). To minimize autofluorescence from the optical fiber, the recording fiber was photobleached using a high-power LED before recording. The day before recording, the mice received an injection of SiR650 via the tail vein.

#### Sucrose

For sucrose delivery, an intraoral cheek fistula was implanted in each mouse. In brief, incisions were made in the cheek and the scalp at the back of the neck. A short, soft silastic tube (inner diameter: 0.5 mm; outer diameter: 1 mm) connected via an L-shaped stainless-steel tube was then inserted into the cheek incision site. The steel tube was routed through the scalp incision, with the opposite end inserted into the oral cavity. After 3 d of recovery from the surgery, the mice were water-restricted for 36 h (until reaching 85% of their initial body weight). The water-restricted, freely moving mice then received 5% sucrose water delivery (approximately 8 μl per trial, with 25-50 trials per session and a trial interval of 20- 30 s).

#### Foot shock

The mice were placed in a shock box and habituated for 30 min. During the experiment, 10 1-s pulses of electricity were delivered at 0.7 mA, with an interval of 90-120 s between trials.

#### Cocaine

Cocaine HCl was obtained from the Qinghai Pharmaceutical Factory and dissolved in 0.9% saline. The mice received an intraperitoneal injection of cocaine (20 mg/kg) in a total volume of 300-400 μl. Photometry signals were recorded for 10-15 min before and 60 min after cocaine injection. The signals were low-pass filtered (0.01 Hz) to remove spontaneous fluctuations in fluorescence.

#### Data analysis of three-color photometry

The photometry data were analyzed using a custom program written in MATLAB. For the sucrose experiment, the baseline was defined as the average fluorescence signal measured for 3-6 s before sucrose delivery; for the foot shock experiment, the baseline was defined as the average fluorescence signal measured for 0-3 s before foot shock delivery; for the cocaine experiment, the baseline was defined as the average fluorescence signal measured for 0-600 s before cocaine injection. To quantify the change in fluorescence across multiple animals, ΔF/F_0_ was normalized using the standard deviation of the baseline signals in order to obtain a *Z*-score.

Signals recorded between adjacent sucrose deliveries (10 s after one sucrose delivery and 5 s before the next sucrose delivery) were used to analyze spontaneous activity (as shown in Fig. 4E1, F1). The DA peaks were identified using the MATLAB *findpeaks* function, with a minimum peak prominence of 2x the standard deviation; standard deviation was calculated based on the baseline following SCH administration. The DA, ACh, and cAMP transients were aligned to the DA peak.

### Immunohistochemistry

Mice were anesthetized and intracardially perfused with PBS followed by 4% paraformaldehyde (PFA) in PBS buffer. The brains were dissected and fixed overnight at 4°C in 4% PFA in PBS. The brains were then dehydrated in 30% sucrose in PBS and sectioned at a thickness of 40 μm using a cryostat microtome (CM1950; Leica). The slices were placed in blocking solution containing 5% (v/v) normal goat serum, 0.1% Triton X- 100, and 2 mM MgCl_2_ in PBS for 1 h at room temperature. The slices were then incubated in AGT solution (0.5% normal goat serum, 0.1% Triton X-100, and 2 mM MgCl_2_ in PBS) containing primary antibodies overnight at 4°C. The following day, the slices were rinsed three times in AGT solution and incubated for 2 h at room temperature with secondary antibodies containing DAPI (5 mg/ml, dilution 1:1,000; catalog no. HY-D0814, MedChemExpress). After three washes in AGT solution, the slices were mounted on slides and imaged using a VS120-S6-W Virtual Slide Microscope (Olympus) equipped with a 10× (NA: 0.4) objective.

Anti-HaloTag primary antibody (rabbit, 1 mg/ml, dilution 1:500; catalog no. G928A, Promega) and iFluor 647-conjugated anti-rabbit secondary antibody (goat, 1 mg/ml, dilution 1:500; catalog no. 16837, AAT Bioquest) were used for HaloDA1.0 and HaloDAmut. Anti-mCherry primary antibody (mouse, 1 mg/ml, dilution 1:1000; catalog no. ab125096, Abcam) and iFluor 555-conjugated anti-mouse secondary antibody (goat, 1 mg/ml, dilution 1:500; catalog no. 16776, AAT Bioquest) were used for jRGECO1a, rACh1h, and ChR2- mcherry. Anti-GFP antibody (chicken, 10 mg/ml, dilution 1:500; catalog no. ab13970, Abcam) and Alexa Fluor 488-conjugated anti-chicken secondary antibody (goat, 2 mg/ml, dilution 1:500; catalog no. ab150169, Abcam) were used for GFlamp2 and ChR2-EYFP.

### Quantification and statistical analysis

Imaging data were processed using ImageJ software (NIH) and custom-written MATLAB (R2020b) programs. Data were plotted using OriginPro 2020b (OriginLab) or Adobe Illustrator CC. The signal-to-noise ratio (SNR) was calculated as the peak response divided by the standard deviation of the baseline fluorescence. Except where indicated otherwise, all summary data are presented as the mean ± s.e.m. All data were assumed to be distributed normally, and equal variances were formally tested. Differences were analyzed using a two-tailed Student’s *t*-test or one-way ANOVA; where applicable, \**P* < 0.05, \*\**P* < 0.01, \*\*\**P* < 0.001, and n.s., not significant (*P* ≥ 0.05).

## ACKNOWLEDGMENTS

We thank Y. Rao for providing the upright confocal microscope. We also thank the optical Imaging platform of the National Center for Protein Sciences at Peking University in Beijing, China, for their support and assistance with the Operetta CLS high-content imaging system and the Nikon A1RSi+ laser scanning microscope. We are grateful to Thinker Tech Nanjing BioScience for their customization and assistance with the fiber photometry system. Some diagrams were created using BioRender.com. We thank the members of the Li lab for helpful suggestions and comments regarding the manuscript.

## AUTHOR CONTRIBUTIONS

Y.L. supervised the project. Y. Zheng and Y.L. designed the study. Y. Zheng developed and optimized the sensors. Y. Zheng performed the experiments related to characterizing the sensors with help from Y. Zhang, G.L., Z.W., Y. Zhuo, F.D., E.J., Y.Y., and K.Z. R.C. performed the confocal imaging of acute brain slices. K.W. performed the zebrafish imaging under the supervision of Y.M. Y. Zheng performed the fiber photometry recording experiments with help from H.D., Y.W., Y.C., J.W., X.M., and S.L. J.Z. performed the chemical conjugation of the HaloTag ligand to chemical dyes under the supervision of Z.C. J.G. and L.L. provided the JF dyes, while K.J. provided other dyes. K.J. and E.S. provided assistance with the chemigenetic strategy. All authors contributed to the interpretation and analysis of the data. Y. Zheng and Y.L. wrote the manuscript with input from all coauthors.

---

**Fig. S1.**
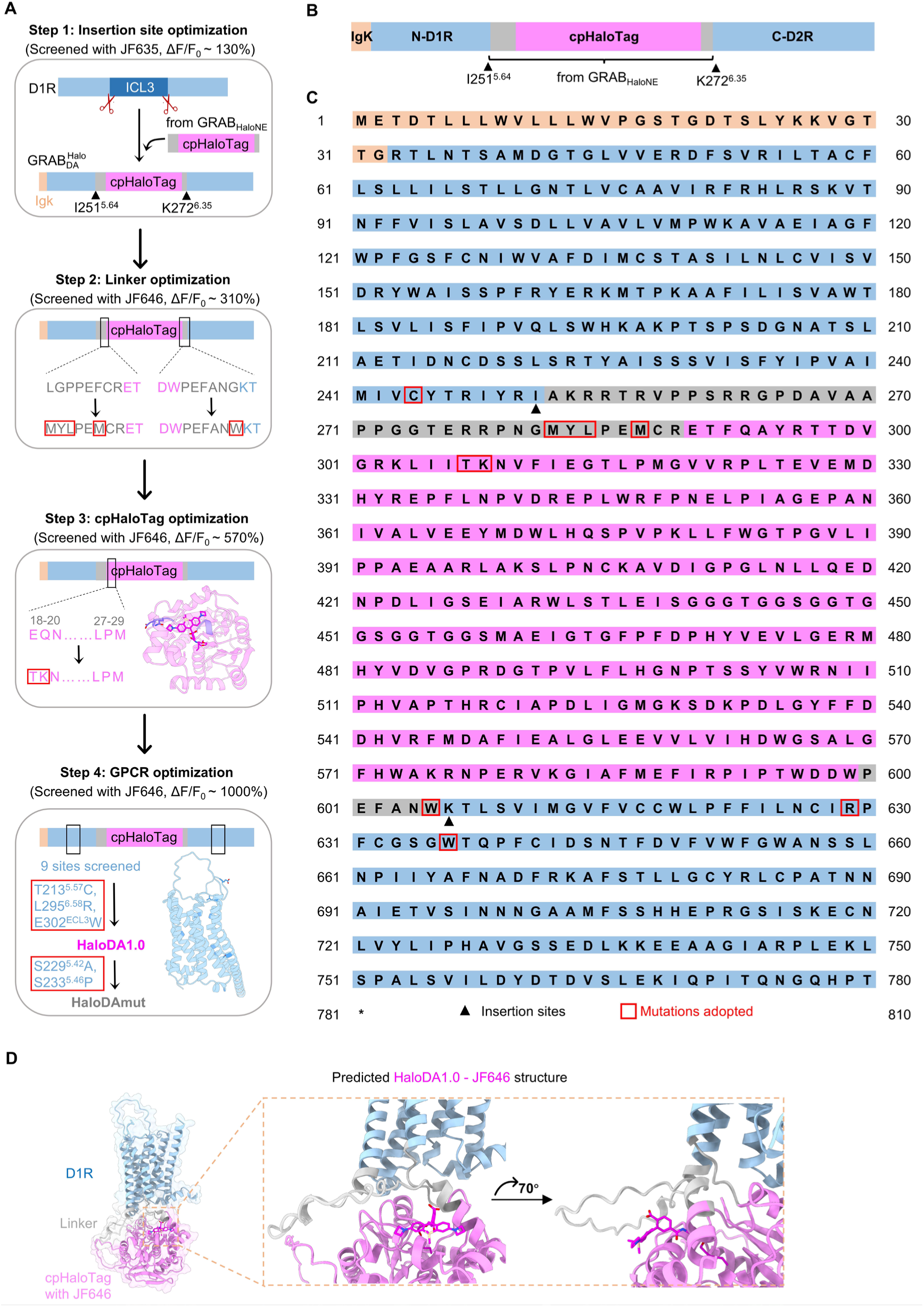
Strategy for optimizing the HaloDA sensor. (**A**) Schematic diagram showing the design and optimization of HaloDA1.0 and HaloDAmut. The structure in step 3 is from the resolved cpHaloTag structure (PDB: 6U32); the structure in step 4 is from the resolved D1R structure (PDB: 7JVQ). IgK refers to the IgK leader sequence. (**B** and **C**) Schematic depiction (**B**) and amino acid sequence (**C**) of HaloDA1.0; the black triangles indicate the insertion sites of cpHaloTag with linkers into D1R, and the red boxes indicate mutation sites introduced during sensor optimization. (**D**) Predicted structure of HaloDA1.0 using AlphaFold 3(*69*). JF646 conjugated with the HaloTag ligand was docked into the structure by alignment with the published cpHaloTag-dye structure (PDB: 6U32).

**Fig. S2.**
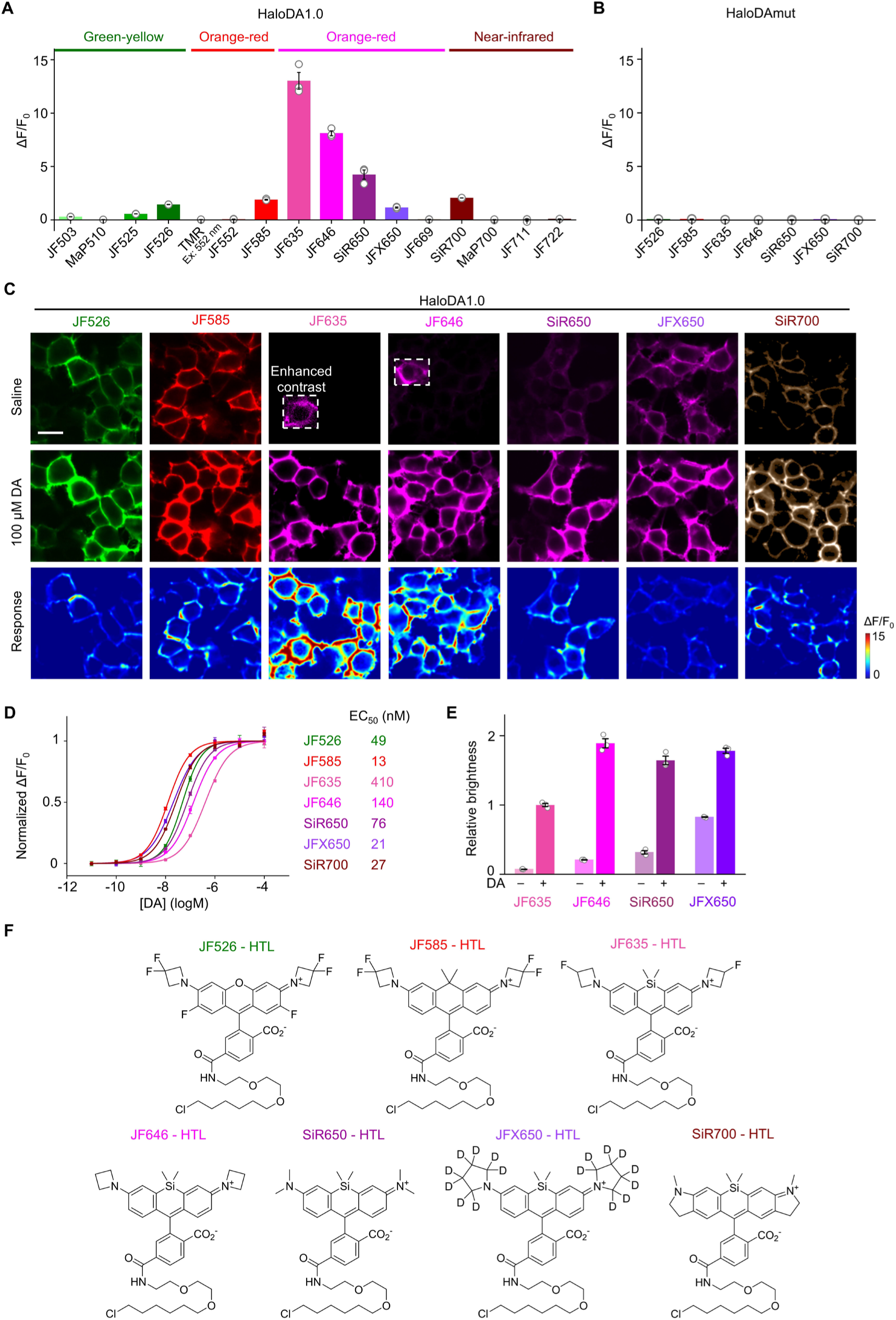
Performance of HaloDA1.0 sensors labeled with various dyes. (**A** and **B**) Maximum ΔF/F_0_ of HaloDA1.0 (**A**) and HaloDAmut (**B**) expressed in HEK293T cells and labeled with the indicated dyes; n = 3 wells with 300–500 cells per well. (**C**) Representative images of HEK293T cells expressing HaloDA1.0 and labeled with the indicated dyes before and after application of 100 μM DA. Scale bar, 20 μm. (**D** and **E**) Normalized dose-response curves (**D**) and relative brightness (**E**, normalized to JF635-labeled HaloDA1.0 measured in the presence DA) of HaloDA1.0 expressed in HEK293T cells and labeled with the indicated dyes; n = 3 wells with 300–500 cells per well. (**F**) Structures of the indicated seven dyes conjugated with the HaloTag ligand (HTL).

**Fig. S3.**
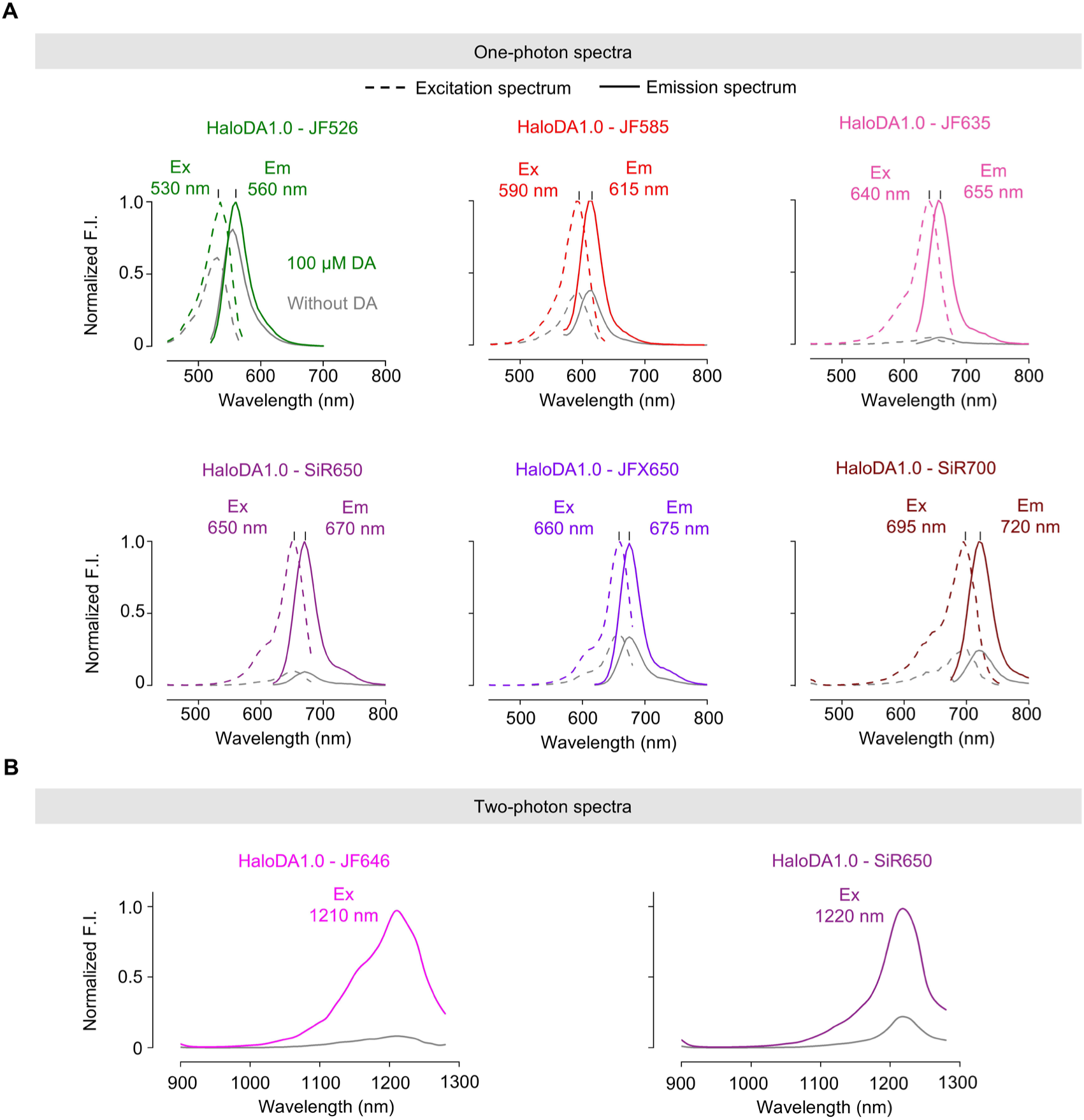
Spectral properties of HaloDA1.0 sensors labeled with various dyes. (**A**) One-photon excitation (Ex, dash line) and emission (Em, solid line) spectra of HaloDA1.0 labeled with the indicated dyes and measured in the absence (gray line) and presence of 100 μM DA (colored line). (B) Two-photon excitation and emission spectra of HaloDA1.0 labeled with the indicated dyes and measured in the absence (gray line) and presence of 100 μM DA (colored line).

**Fig. S4.**
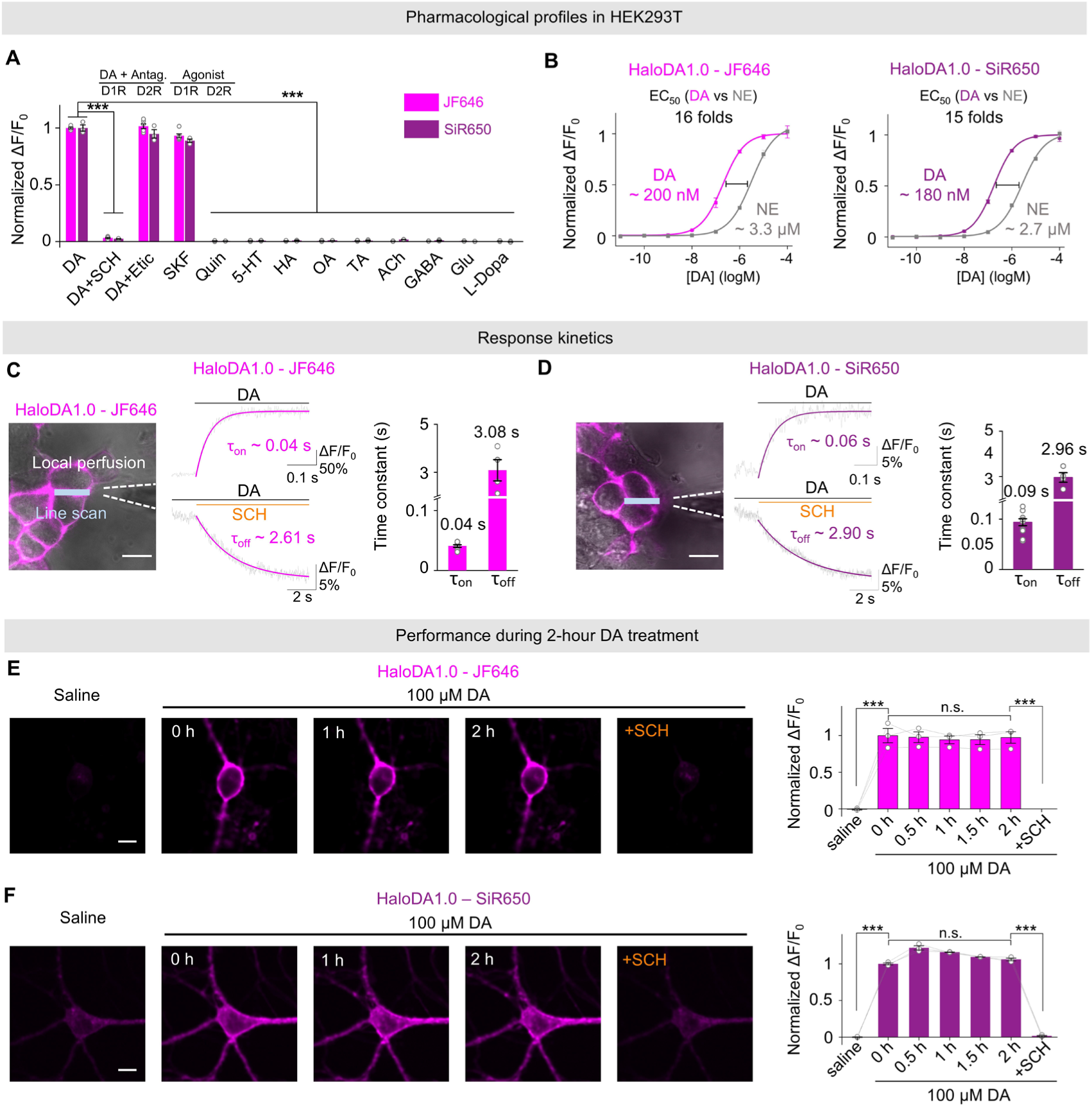
Characterization of the HaloDA1.0 sensor expressed in cultured cells. (**A**) Summary of the response of HaloDA1.0 expressed in cultured HEK293T cells and labeled with JF646 or SiR650. All chemicals were applied at 1 μM; n = 3 wells for each condition. (**B**) Dose-response curves of HaloDA1.0 expressed in cultured HEK293T and labeled with JF646 (left) or SiR650 (right), in response to DA or NE; n = 3 wells for each condition. (**C** and **D**) Schematic illustration (left), representative traces (middle), and group summary (right) of the response to locally puffing DA or SCH in order to measure the kinetics of HaloDA1.0 labeled with JF646 (**C**) or SiR650 (**D**). τ_on_ was measured following a puff of DA, while τ_off_ was measured following a puff of SCH in the presence of DA; n = 4–9 cells each. Each trace was fitted with a single-exponential function. Scale bars, 20 μm. (**E and F**) Representative images (left) and group summary of normalized ΔF/F_0_ (right) measured in cultured neurons expressing HaloDA1.0 and labeled with JF646 (**E**) or SiR650 (**F**) before and up to 2 hours after application of 100 μM DA, followed by the addition of 100 μM SCH; n = 3 coverslips for each condition. Scale bars, 10 μm.

**Fig. S5.**
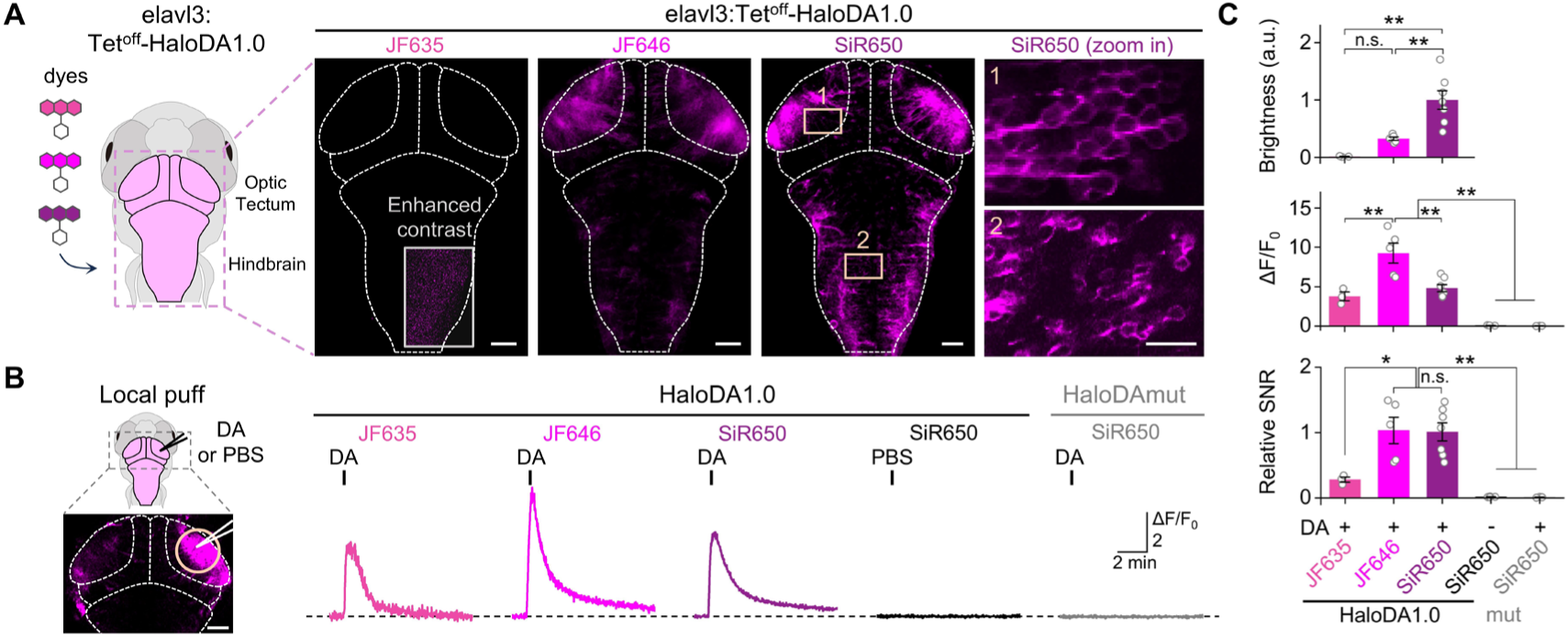
Performance of HaloDA1.0 sensors in zebrafish labeled with various dyes. (**A**) (Left) Schematic diagram of the zebrafish larvae’s head with various dye labeling and the indicated field of view for confocal imaging. (Right) Representative images of the expression of HaloDA1.0 labeled with JF635, JF646, or SiR650. Scale bars, 50 μm. Two expanded views showing single-cell resolution in the indicated brain regions in SiR650- labeled zebrafish are shown on the right (scale bar, 20 μm). (**B**) (Left) Schematic diagram and representative image of a local puff of DA or PBS onto the zebrafish brain. The orange circle (100 μm diameter) indicates the ROI used for further analysis. Scale bar, 50 μm. (Right) Representative traces of the change in HaloDA1.0 or HaloDAmut fluorescence measured under the indicated conditions. The short vertical black lines indicate local puffs. (**C**) Group summary of the brightness, ΔF/F_0_, and relative SNR in response to local puff under the indicated conditions; n = 3-7 zebrafish per group.

**Fig. S6.**
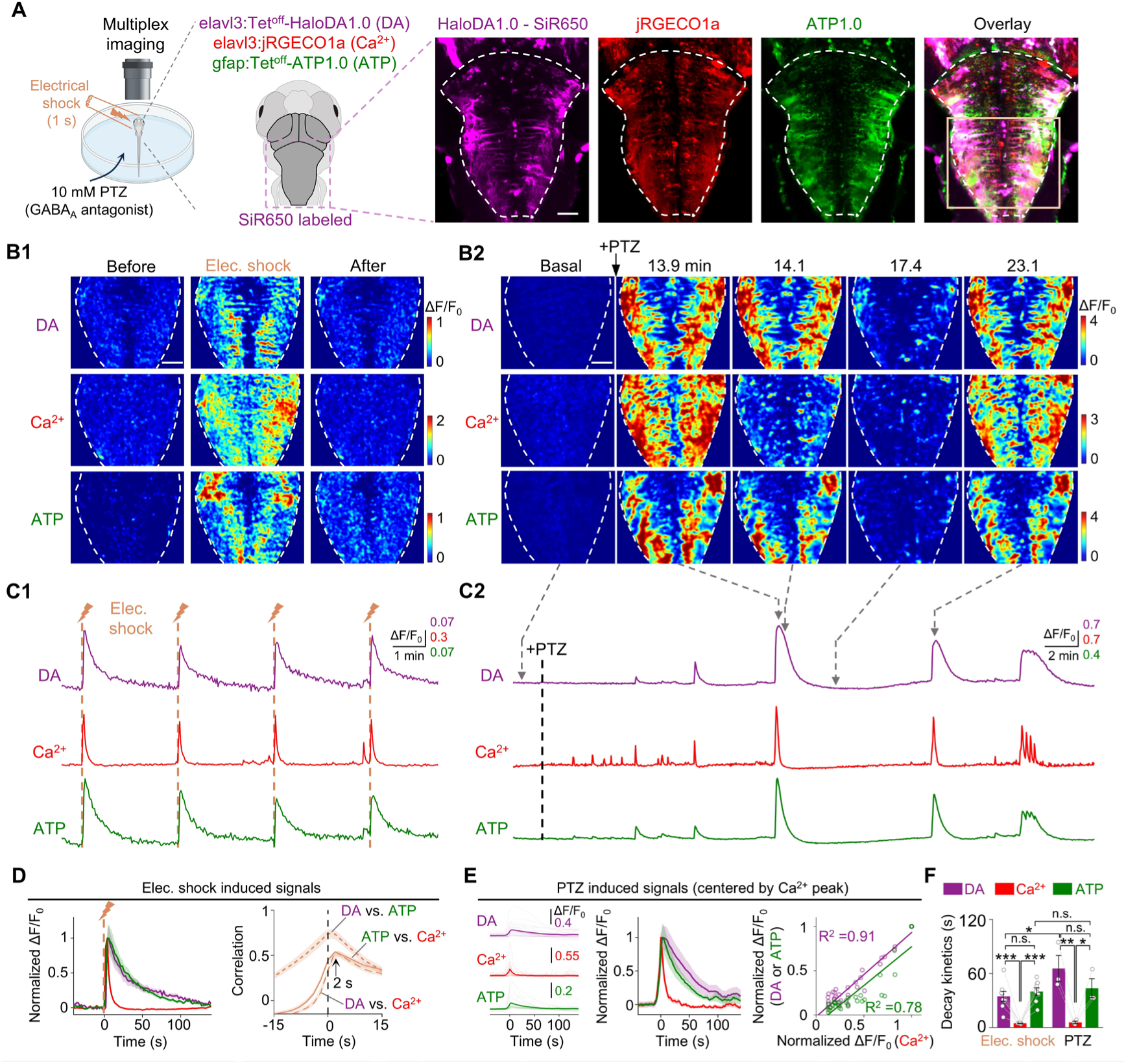
Multiplex imaging in zebrafish. (**A**) Schematic diagram and representative images of multiplex imaging in the hindbrain of zebrafish in response to a 1-s electrical shock or 10 mM pentylenetetrazole (PTZ) application. The zebrafish were labeled with SiR650. The orange box in the overlay indicates the ROI used for further analysis. Scale bar, 50 μm. (**B** and **C**) Pseudocolor images (**B**) and example traces (**C**) of DA, Ca^2+^, and ATP signals measured during electrical shock **(B1** and **C1)** and PTZ application **(B2** and **C2)**. Scale bars, 50 μm. (**D**) Normalized fluorescence response (left) and cross-correlation (right) of the indicated pairs of sensor signals measured during electrical shock; n = 8 zebrafish. (**E**) DA, Ca^2+^, and ATP signals measured during PTZ application. (Left) Peak fluorescence responses obtained by centering all three sensor signals with the peak Ca^2+^ signal. (Middle) Normalized fluorescence response of all three signals. (Right) Scatter plot of the normalized peak amplitude of all three signals. Individual peak amplitude was normalized to the maximum peak amplitude for each sensor signal. The magenta circles indicate the correlation between DA and Ca^2+^, while the green circles indicate the correlation between ATP and Ca^2+^. The data were fitted with a linear function. A total of 33 peaks were selected in 3 zebrafish. (**F**) Group summary of the decay kinetics of all three sensor signals measured during electrical shock (n = 8 zebrafish) or PTZ application (n = 3 zebrafish). The values were obtained by fitting the traces with a single-exponential function.

**Fig. S7.**
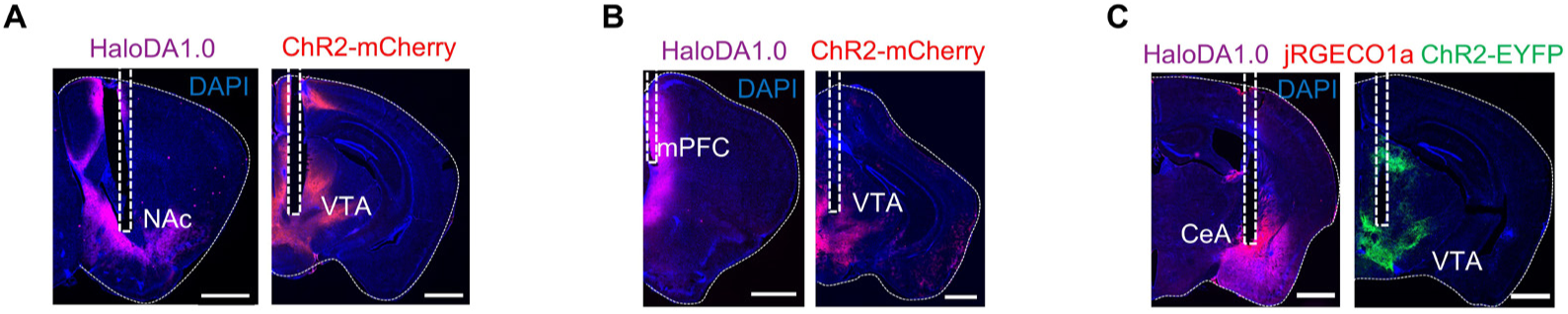
Validation of optogenetic expression in mice. Histological verification of the expression of the indicated sensors and optogenetic actuators in the VTA and NAc (**A**), VTA and mPFC (**B**), and VTA and CeA (**C**). The dashed lines indicate the location of the optical tract. Scale bars, 1 mm.

**Fig. S8.**
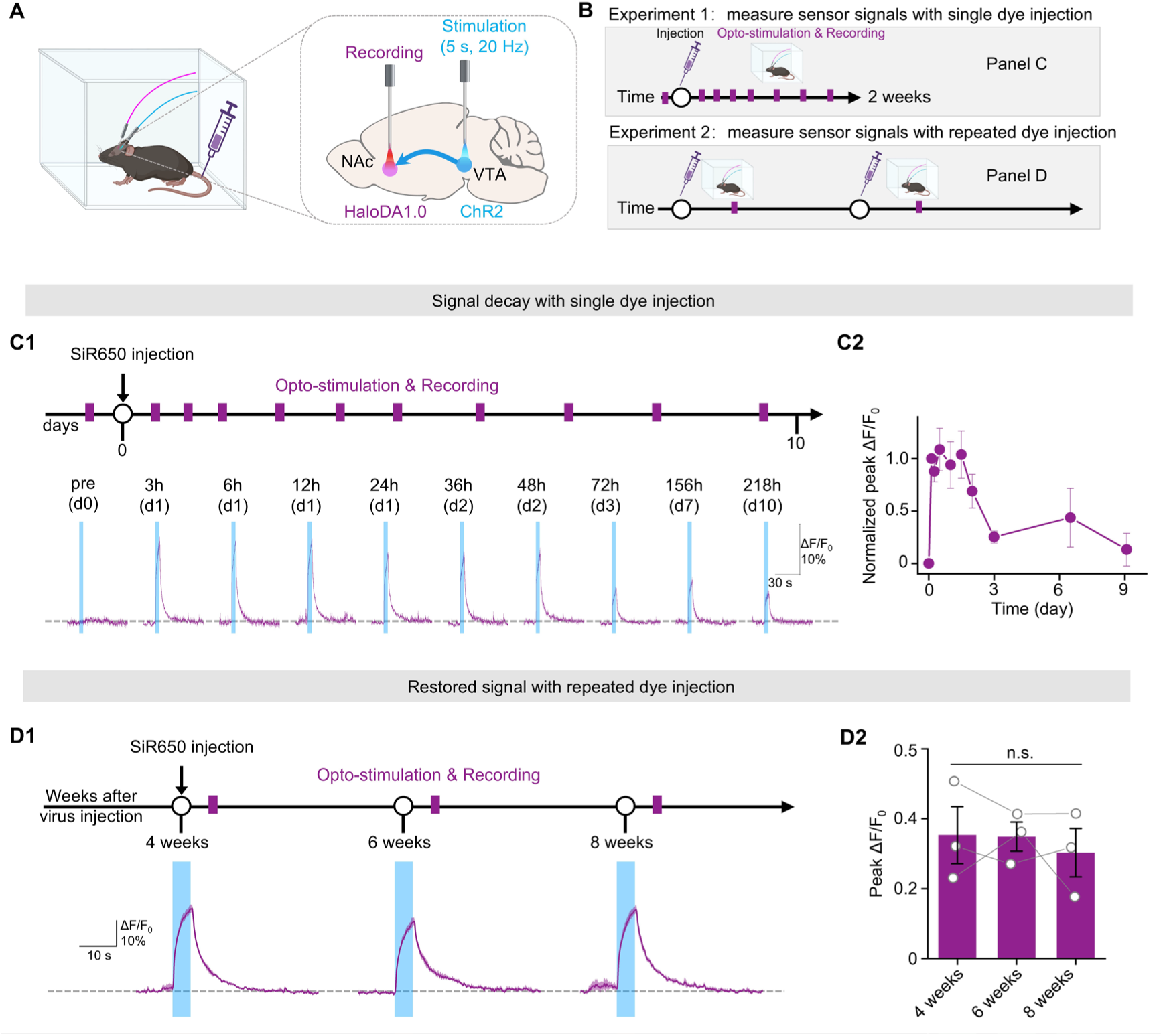
Measuring sensor signals after a single or repeated dye injections. (**A**) Schematic diagram depicting the strategy for fiber photometry recording of HaloDA1.0 in the NAc upon optogenetic stimulation of VTA neurons. (**B**) Schematic diagram depicting the experimental protocol for measuring sensor signals, with a single injection of 100 nmol SiR650 in the tail vein (experiment 1, top) or repeated injections of 100 nmol SiR650 (experiment 2, bottom). (**C**) Representative fluorescence responses to optogenetic stimuli (**C1**) and group summary of normalized peak ΔF/F_0_ (**C2**) measured before dye injection and at the indicated time points after a single injection of dye; n = 3 mice. The vertical blue shading indicates the optogenetic stimuli. (**D**) Representative fluorescence responses to optogenetic stimuli (**D1**) and group summary of peak ΔF/F_0_ (**D2**) measured with repeated dye injections in weeks 4, 6, and 8. Each measurement was performed 12 hours after dye injection; n = 3 mice.

**Fig. S9.**
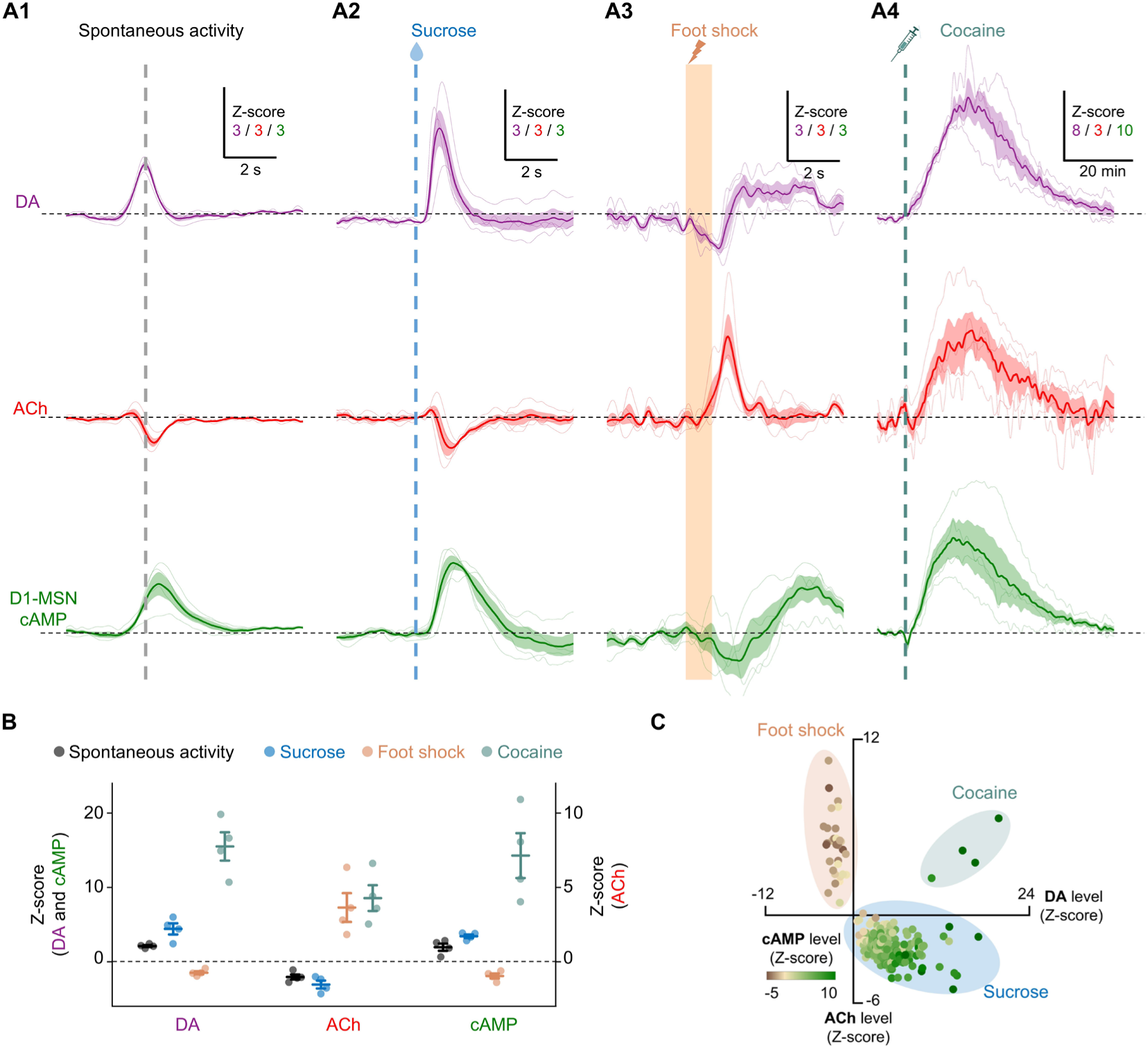
Measuring the fluorescence responses of DA, ACh, and cAMP in vivo. (**A**) The change in fluorescence for the HaloDA1.0 (DA), rACh1h (ACh), and DIO-GFlamp2 (D1-MSN cAMP) sensors measured during spontaneous activity (**A1**) and in response to sucrose (**A2**), foot shock (**A3**), and cocaine application (**A4**). The thin traces represent the fluorescence changes measured in an individual mouse, while the thick traces indicate the average fluorescence change; n = 4 mice for each condition. (**B**) Group summary of the peak or trough responses for all three sensor signals under the indicated conditions; n = 4 mice. (**C**) Scatter plot of the peak/trough amplitude of the three sensor signals measured under the indicated conditions; n = 4 mice. Each point represents an individual trial. The ACh response is plotted on the *y*-axis, the DA response is plotted on the *x*-axis, and the color of each data point indicates the cAMP response.

**Fig. S10.**
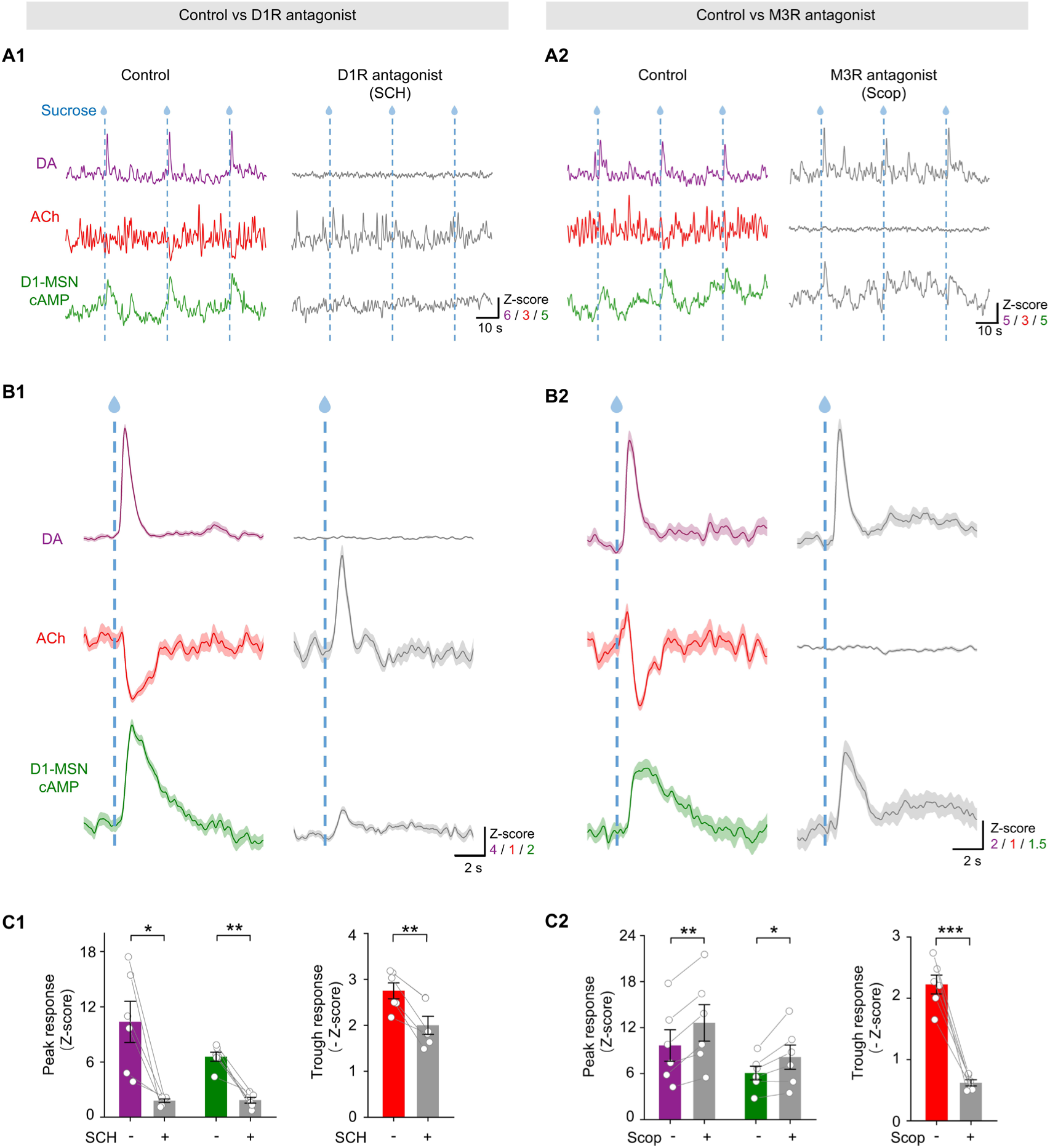
Pharmacologic validation during three-color recording. Representative traces of the change in fluorescence (**A**), average traces (**B**), and group summary of peak *Z*-scores (**C**) measured for DA, ACh and D1-MSN cAMP sensors under control conditions and following an i.p. injection of 8 mg/kg SCH (**A1**, **B1**, and **C1**) or 10 mg/kg Scop (**A2**, **B2**, and **C2)**.

